# Epigenetic reprogramming by promoter hypomethylation is linked to aberrant gene expression and transcriptomic plasticity in colorectal cancer

**DOI:** 10.64898/2026.11.04.763625

**Authors:** Luís Nunes, Ina A. Eilertsen, Rita Pinto, Hege M. Vedeld, Ingrid M. Bolstad, Usman Saeed, Ole H. Sjo, Sheraz Yaqub, Arild Nesbakken, Kushtrim Kryeziu, Guro E. Lind, Ragnhild A. Lothe, Anita Sveen

## Abstract

Colorectal cancer (CRC) has high phenotypic plasticity, but regulatory mechanisms of aberrant cell states are not well described. We investigated DNA methylation-based regulation of cancer cell-intrinsic transcriptomic heterogeneity in 1,183 multiregional primary tumor samples, liver metastases and tumor organoids. Most genes with strong methylation-transcriptomic association reflected canonical promoter hypermethylation and expression repression related to the CpG island methylator phenotype (CIMP). However, promoter hypomethylation and overexpression of *APOLD1* and *ELF5* identified two largely non-overlapping subgroups of canonical CIMP-negative CRCs enriched with stem-like and transit-amplifying cell states. *APOLD1* tumors were further highly proliferative while *ELF5* tumors were slow-cycling and associated with good survival in early-stage disease. Target genes showed patient-specific expression changes in matched primary-metastasis samples, and *ELF5* aberrant metastases showed *ex vivo* sensitivity to putative targeting of stem cells by PI3K/mTOR inhibition, fetal de-differentiation and transition into non-canonical intestinal states. This implicates focal promoter hypomethylation in phenotypic plasticity in CRC metastasis.

## Background

Phenotypic plasticity allows cancer cells to rapidly adapt to developmental, microenvironmental, and therapeutic pressures, resulting in transcriptomic states that can sustain tumor progression, metastatic competence, and treatment resistance^1^. Colorectal cancer (CRC) is a highly plastic cancer type. The malignant cells exist in a continuum of transcriptomic states and cell state transitions result in progressive plasticity during metastasis^2,3^. Furthermore, drug-tolerant states adopted by CRC cells under treatment pressure are reversible^4^, highlighting the dynamics of phenotypic plasticity and pointing to epigenetic regulatory mechanisms. Integrated genomic and transcriptomic analyses of individual tumor gland has further shown that most intratumoral variation of gene expression in CRC cannot be explained by genetic ancestry^5^, and it is important to obtain a better understanding of the epigenetic mechanisms governing phenotypic plasticity.

Epigenetic mechanisms can establish and maintain transcriptomic states and are both heritable and reversible through cell division, providing dynamic transcriptomic regulation^6^. Studies of epigenomic and genomic co-evolution in CRC have shown that regulation of chromatin accessibility contributes to tumor development^7^, and there are widespread associations between aberrant DNA methylation and gene expression^8^. DNA methylation has mostly been studied in the context of hypermethylation of CpG-islands in promoter regions, with subsequent gene expression silencing of targeted genes. The best-established example is the CpG island methylator phenotype (CIMP), which is characterized by coordinated promoter hypermethylation, including the DNA mismatch repair gene *MLH1*. This results in microsatellite instability (MSI) and a high tumor mutational burden. CIMP+ tumors are further associated with *BRAF* mutation, proximal tumor location, old patient age, and female sex^9–11^. DNA hypomethylation has received considerably less attention, but a few specific events have been shown to contribute to oncogenesis through gene reactivation. One of the earliest examples is hypomethylation of the *HRAS* proto-oncogene^12^ and a well-established example in CRC is promoter demethylation and increased expression of *EREG* during the adenoma-to-carcinoma transition, associated with enhanced *EGFR* signaling and response to EGFR targeted therapy^13^. Beyond promoter regions, cancer-associated DNA methylation changes can occur at enhancers, open chromatin regions, and other regulatory elements that influence tissue- and cell-state specific transcription^14^. For example, hypomethylation of an intronic LINE-1 transposon element has been associated with increased expression of the *MET* proto-oncogene in CRC metastasis^15^. However, it remains unclear whether such changes are associated with transcriptomic states beyond the characteristics of CIMP, and how they shape spatiotemporal tumor heterogeneity.

Addressing this question in bulk tumor material containing a mixture of cell types is challenging because both DNA methylation and gene expression are strongly tissue- and cell-type specific. Regional differences in the composition of the tumor microenvironment can therefore obscure cancer cell-intrinsic regulatory mechanisms. Single-cell profiling has the potential to resolve this issue, but remains expensive and technically challenging, particularly for DNA methylation analysis^16,17^. We previously showed that bulk transcriptomics of spatially distinct regions of primary CRCs can identify cancer cell-intrinsic features based on genes with low intra-tumor expression variation^18^. Here, we extend this framework by integrating DNA methylation profiling of multiregional tumor samples to identify cancer cell-intrinsic methylation-expression patterns and epigenetic marks of distinct cancer cell states and phenotypic plasticity in the context of spatiotemporal tumor heterogeneity in CRC metastasis.

## Results

### Concordant CIMP status among multiregional samples of primary CRCs

Tumor heterogeneity of the DNA methylation landscape in CRC was initially analyzed in 2-4 spatially distinct regions from each of 94 primary tumors, including CIMP status, *MLH1* promoter methylation, and epigenome-wide profiling on Illumina Infinium EPIC arrays (*n* = 92 patients and 274 samples after quality control; Figure S1). The samples have previously been analyzed for MSI status, *BRAF* and RAS (*KRAS* and *NRAS)* hotspot mutations, and transcriptomics^18^. CIMP status was highly concordant among multiregional samples in each tumor (Figure 1A). Only one tumor showed phenotype discordance, with one CIMP+ and two CIMP-samples distinguished by only one of the markers in the Weisenberger panel (*RUNX3*; Figure S2)^19^. Two synchronous tumors from each of two patients were both either CIMP+ or CIMP-. CIMP+ tumors had frequent *MLH1* promoter hypermethylation and MSI (20/29 non-synchronous tumors with homogeneous CIMP+; 69%; *P* = 5×10^-12^ by Fisher’s exact test; Table S1). All CIMP+ tumors were also mutated for either *BRAF* V600E or RAS hotspots. However, only *BRAF* V600E mutations were enriched among CIMP+ tumors (83% vs 3%), while the prevalence of RAS mutations was higher in the CIMP-group (48% vs 17%; Fisher *P* = 2×10^-15^). CIMP+ tumors were more often located in the right side of the colon (90% CIMP+ vs 22% CIMP-; Fisher *P* = 3×10^-9^) and were associated with older patient age (48% vs 23% ≥75 years; Fisher *P* = 0.01) and female sex (72% vs 38%; Fisher *P* = 0.003; Table S1), consistent with prior epidemiological studies^10,11,20^. CIMP+ tumors were also frequently diagnosed at advanced cancer stages, but this association was not statistically significant (Fisher *P* = 0.09).

**Figure 1.**
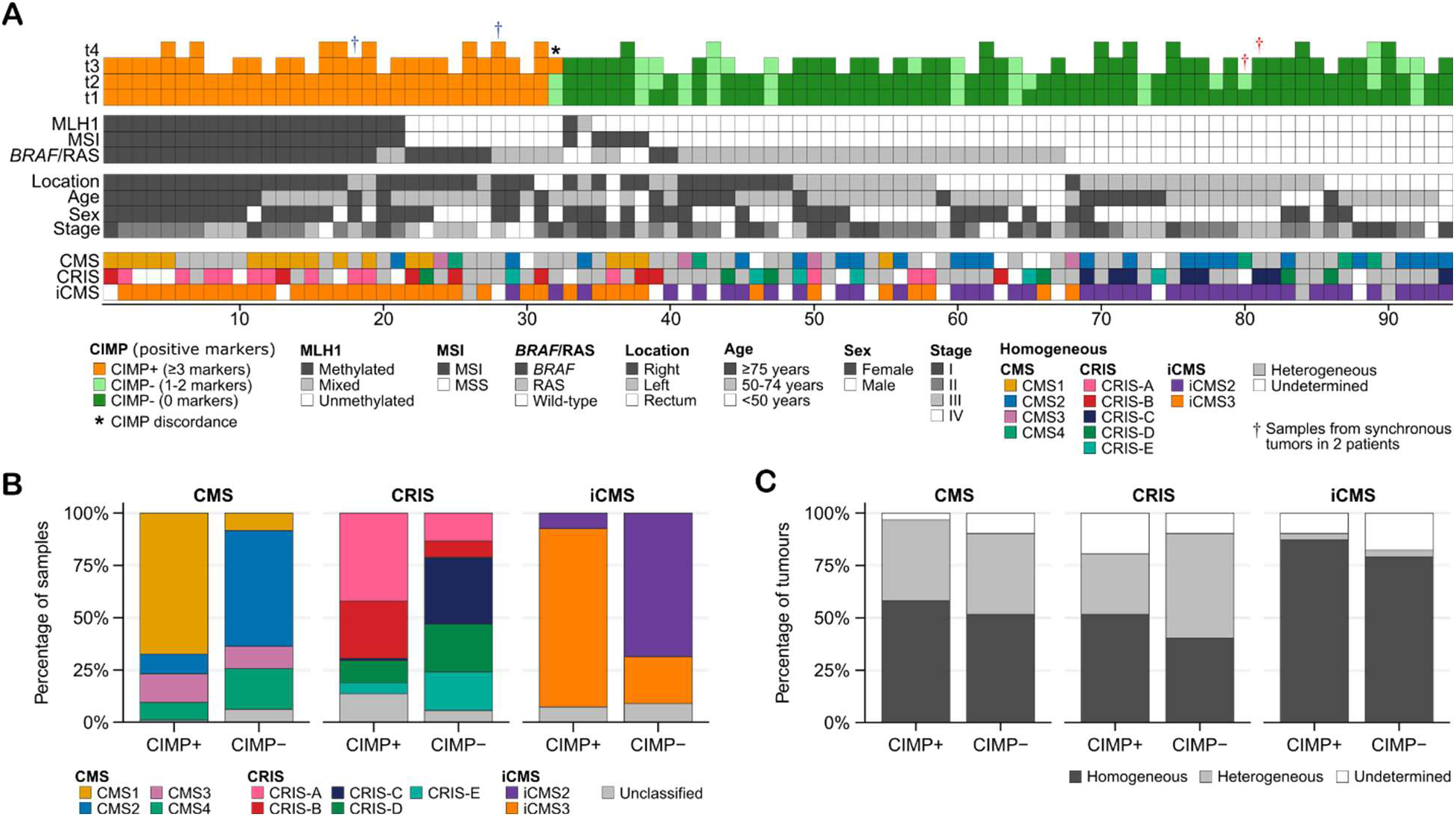
Multiregional landscape of CpG island methylator phenotype (CIMP) status in primary colorectal cancer (CRC). **(A)** CIMP classification of 2-4 spatially distinct regions from each of 94 primary CRCs (total *n* = 274 samples), ordered by clinicopathological and tumor characteristics. Each column represents one tumor. Multiregional samples (t1–t4) are color-coded according to the number of methylated loci based on the Weisenberger five-marker panel. Molecular annotation bars below indicate *MLH1* promoter methylation status, MSI status, hotspot mutations in *BRAF*, *KRAS*, and *NRAS* (RAS), as well as transcriptomics classifications (CMS, CRIS, and iCMS) per tumor. Discordant molecular classifications among multiregional samples are marked as heterogenous or undetermined. **(B)** Stacked bar plots showing the distribution of transcriptomic subtypes stratified by CIMP status, colored by sample-level classification. **(C)** Stacked bar plots showing transcriptomic subtype distributions by CIMP status, colored according to intra-tumor heterogeneity of subtype classification.

There was substantial intra-tumor heterogeneity of transcriptomic classifications according to the consensus molecular subtypes (CMS) and CRC intrinsic subtypes (CRIS) among the multiregional samples^18^, but this did not vary according to CIMP status, and CIMP+ tumors were enriched with CMS1 and CRIS-A/B (Figure 1B-C; Table S1). Furthermore, all CIMP+ tumors with homogeneous classification according to the intrinsic CMS (iCMS) were iCMS3, except for a single CIMP+ iCMS2 tumor (iCMS3: 86% CIMP+ vs 20% CIMP-; Fisher *P* = 2×10^-9^). These findings indicate particularly strong epigenetic support for the cancer cell-intrinsic transcriptomic classification.

### Annotation of active promoters regulated by DNA methylation in CRC

Before conducting epigenome-wide methylation analyses, we re-annotated all Illumina EPIC array^21^ probes using regulatory-element information from the ENCODE project^22^. This was to address the limitations of the default probe annotation, which is largely gene-centric and can overestimate promoter-associated probes and underrepresent other regulatory elements, including enhancers. Probes were mapped to both cell type-agnostic and large intestine-specific candidate cis-regulatory elements retrieved through SCREEN, the ENCODE web interface that integrates regulatory annotations across multiple tissue and cell types and cancer cell lines (see Methods; Table S2)^22^.

Using the default Illumina annotation, 49% of the probes were assigned to promoter regions and this proportion increased to 61.8% when probes with dual promoter-enhancer annotations were also included. In contrast, ENCODE annotated only 14.1% and 10.2% of the probes as promoter-like elements in the cell type-agnostic and large intestine-specific regulatory maps, respectively. The reverse pattern was observed for enhancer-like elements, and 9.7% of probes were annotated as enhancers using the default Illumina annotation, compared with 45.9% using the ENCODE cell type-agnostic map. Notably, only 10% of probes overlapped enhancer-like elements in the large intestine-specific map, consistent with the strong cell type-specificity of enhancer activity. Accordingly, most probes in the large intestine-specific annotation mapped to repetitive, gene body or intergenic regions (Figures 2A and S3), reflecting that only a subset of potential promoter and enhancer elements is active or marked as regulatory elements in a specific tissue context.

**Figure 2.**
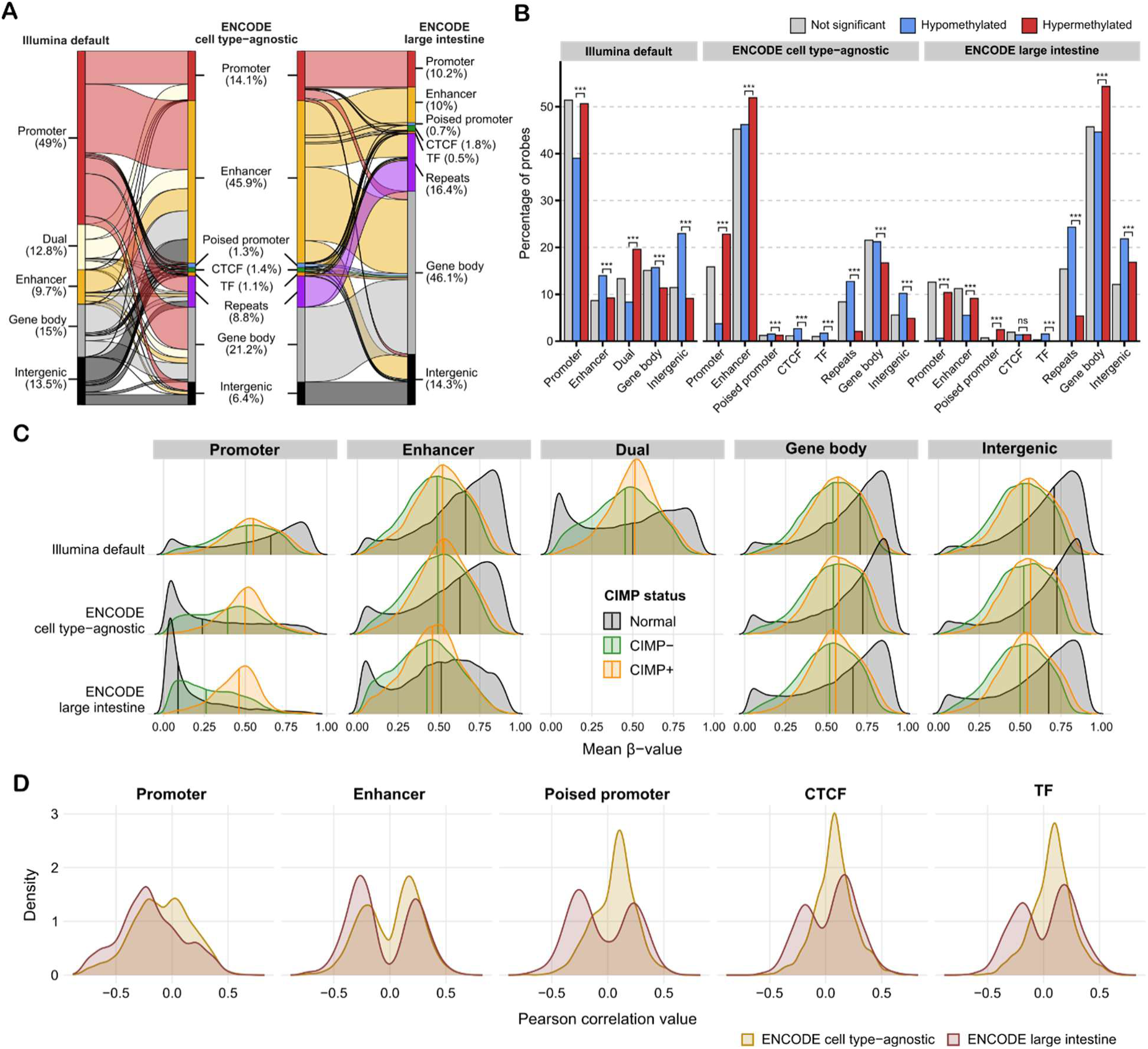
Reannotation of Illumina EPIC array methylation probes using ENCODE data. **(A)** Alluvial diagrams comparing Illumina default probe annotation with ENCODE-derived annotations, including cell-type agnostic and large intestine-specific categories. In the Illumina classification, “dual” denotes probes overlapping both promoter and enhancer regions. Annotations of repeats in the ENCODE-based classifications were obtained from RepeatMasker. **(B)** Bar plots of the percentage of probes with hypomethylation, hypermethylation, or no difference in methylation levels between tumors and healthy colonic mucosa, grouped by probe annotation category. Stars indicate FDR-adjusted chi-square significance (ns: not significant, *FDR < 0.05, **FDR < 0.01, ***FDR < 0.001) between hypo-and hypermethylated probes. **(C)** Density ridge plots of mean β-values stratified by probe annotation category, colored by CIMP status for tumor samples and including normal samples for reference. Vertical black lines indicate median values. **(D)** Density ridge plots of Pearson correlation coefficients between probe methylation β-values and predicted target gene expression, grouped by probe annotation category according to ENCODE and colored according to tissue-specificity of the annotation.

Differential methylation analysis between tumor and healthy colonic tissue samples was used to evaluate cancer-specificity of the candidate regulatory elements (*n* = 274 and 19 samples, respectively; Figure S4; Table S3). Probes hypermethylated in tumors were strongly enriched at promoter-like elements in the ENCODE cell type-agnostic and large intestine-specific annotations (odds ratio, OR = 7.6 and 18.3, respectively, relative to hypomethylated probes; FDR *P* < 0.001), while enrichment at enhancer-like elements was more modest (OR = 1.3 and 1.7, respectively; FDR *P* < 0.001). Using the default Illumina annotation, promoter enrichment was weaker (OR = 1.6; FDR *P* < 0.001) and enhancers were depleted of hypermethylated probes (OR = 0.6; FDR *P* < 0.001; Figure 2B; Table S4). Thus, compared with the default Illumina annotation, ENCODE-based annotations more clearly captured the expected preferential accumulation of hypermethylation at promoter regions in tumors, and showed higher methylation β-values at promoter regions in tumors than healthy tissue samples (Figure 2C). Stratification by CIMP status confirmed that methylation was generally higher in CIMP+ tumors for all regulatory elements, and the difference between CIMP+ and CIMP-tumors was largest for promoters according to ENCODE annotation, particularly with the large intestine-specific annotation (Figures 2C and S5).

Differences between the cell type-agnostic and large intestine-specific annotations from ENCODE were further analyzed based on associations with target gene expression. Promoter-like elements in the large intestine-specific annotation showed stronger inverse correlations between methylation β-values and target gene expression across tumors than the corresponding elements in the cell type-agnostic annotation (Figure 2D). These data support the use of tissue-specific regulatory annotations to identify methylation-expression associations with greater functional relevance in CRC. Notably, enhancers, poised promoters, and CTCF and transcription factor binding sites in the large intestine-specific annotation showed bimodal correlation distributions consistent with a more context-dependent methylation-expression relationship at these regulatory elements (Figure 2D).

### Low intra-tumor heterogeneity of DNA methylation at core regulatory elements of cancer cells

To focus on cancer cell-intrinsic features, we leveraged our previous transcriptomics study showing that intra-tumor heterogeneity is largely driven by variation in the tumor microenvironment and cell type composition, whereas cancer cell-intrinsic signals can be captured by features with low intra-tumor heterogeneity among multiregional samples^18^. To determine whether this principle also applied to DNA methylation, we followed a similar approach to score probes with aberrant, cancer-specific DNA methylation (Table S3) into three categories of intra-tumor heterogeneity based on the variation of β-values within tumors versus among tumors in the multiregional samples set (ITH-low, ITH-intermediate, and ITH-high; see Methods). This approach identified 80 ITH-high (0.05% of the scorable probes), 66,315 ITH-intermediate (44.1%), and 83,869 ITH-low (55.8%) methylation probes (Figure S6; Tables S5 and S6). ITH-low probes were more frequently mapped to promoter-like elements in the ENCODE cell type-agnostic and large intestine-specific maps (OR = 2.8 and 8.5, respectively; FDR *P* < 0.001) and CpG islands (OR = 6.6; FDR *P* < 0.001) and showed higher methylation levels (OR = 3.2; FDR *P* < 0.001) than ITH-high or ITH-intermediate probes (Figure S7A). These findings indicate that ITH-low methylation probes are enriched at core regulatory elements. Only regulatory elements represented by at least three ITH-low probes were retained for downstream analyses (Figure S7B), and most of these were annotated as promoters (59%) or enhancers (33%) in both the cell type-agnostic and large intestine-specific regulatory maps from ENCODE (Figure S7C).

Principal component analysis (PCA) based on ITH-low methylation features showed that CIMP status was a strong driver of variance and separated the tumor samples along PC1 (correlation ratio η = 0.81; Figure 3A-B). In contrast, PCA based on ITH-high/intermediate probes correlated poorly with CIMP status, and PC1 was strongly correlated with the estimated tumor purity of the samples (Spearman’ p = 0.75; Figure S8A-B). To further interpret the biological variation captured by the methylation components, we correlated the PC scores with transcriptomic single-sample gene set enrichment scores from a curated collection (*n* = 157 gene sets; Table S7). PC1 from ITH-low methylation features correlated most strongly with cancer cell-intrinsic transcriptomic programs (Figure 3C), whereas PC1 from ITH-high/intermediate features correlated most strongly with stromal and mesenchymal-like signals (Figure S8C). These data support that ITH-low methylation probes from multiregional bulk tumor samples preferentially capture cancer cell-intrinsic variation.

**Figure 3.**
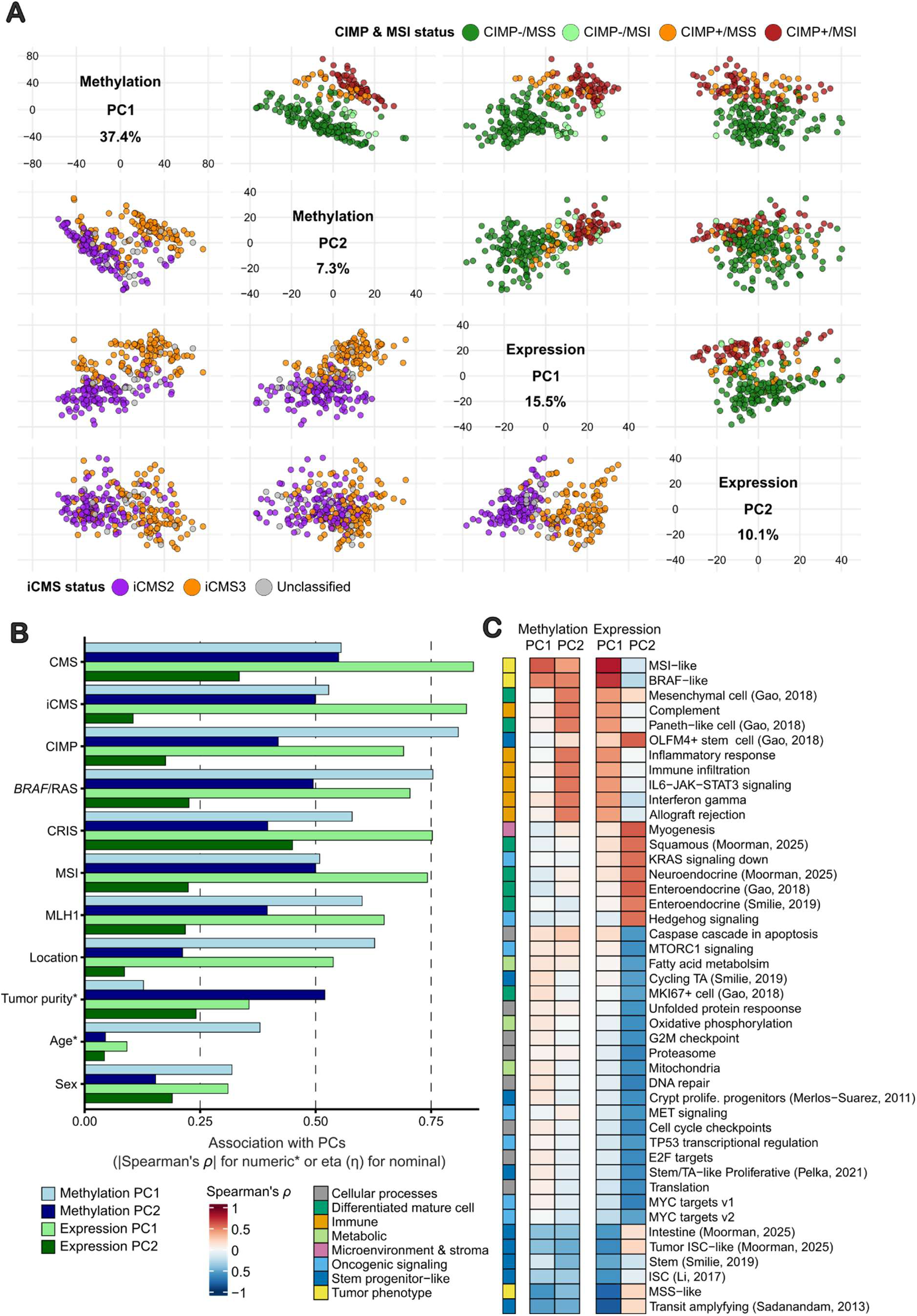
Comparison of ITH-low expression and methylation data. **(A)** Scatter plot matrix (SPLOM) of principal component analysis (PCA) results (PC1 and PC2) for each dataset. For methylation, PCA was performed on M-values restricted to ENCODE cCREs with at least three ITH-low probes annotated, using the median β-value per cCRE. Upper panels are colored by combined CIMP and MSI status, whereas lower panels are colored by iCMS subtype classification. Percentages indicate the variance explained ratio by each PC in the respective dataset. **(B)** Associations between PCs with molecular and clinicopathological characteristics. Associations with continuous variables were assessed using Spearman correlation, and associations with categorical variables using correlation ratio eta (η). Bars are colored by PC. **(C)** Heatmap of gene set variation analysis scores, showing correlations between gene set activity and PCs. Color indicates Spearman correlation coefficients. Only gene sets with at least one absolute correlation ≥ 0.5 are shown.

The methylation-based ITH-low PC1 and PC2 were also strongly correlated with the transcriptomic ITH-low PC1 (Spearman’ p = 0.61 and 0.58, respectively, FDR *P* values < 0.01), reflecting strong methylation-based regulation of cancer cell-intrinsic gene expression. These principal components effectively separated stem-like tumors from more differentiated tumors with inflammatory and immune responses, as evaluated by correlation analyses with single-sample enrichment scores (Figure 3C; Table S7). In contrast, transcriptomic PC2 showed only a weak or no correlation with methylation-based PC1 and PC2 (Spearman’ p = -0.21 and -0.06, FDR *P* value < 0.01 and 0.45, respectively). Transcriptomic PC2 was also associated with a distinct set of cellular differentiation, trans-differentiation, and oncogenic signaling programs (Figure 3C), indicating a complementary and additional axis of transcriptomic plasticity that is only weakly reflected in global DNA methylation patterns.

### Aberrant DNA methylation is associated with cancer cell-intrinsic transcriptomic plasticity

Methylation-expression links between putative regulatory elements and their predicted target genes (predictions retrieved from the SCREEN database)^23–26^ were evaluated based on correlation between the methylation β-values and corresponding gene expression levels. These analyses were focused on cancer cell-intrinsic and ITH-low signals, to reduce the potentially confounding effect of the tumor microenvironment (*n* = 1,562 ITH-low methylation-expression pairs, including 801 unique genes). In total, 11.3% (*n* = 177) of the ITH-low pairs showed strong correlation (absolute Pearson correlation coefficient |r| > 0.5 and *P* values < 0.05), the majority of which were negative correlations (*n* = 152, 85.9%; Table S8). In 41 pairs, methylation accounted for a large proportion of gene expression variation (marginal R^2^ > 0.5). Of these, 29 were designated principal methylation-expression correlations, representing the top-ranked correlations by marginal R^2^ for each regulatory element and each gene (Figure 4A and S9). The remaining 12 pairs were classified as secondary correlations (Figure S10). All principal correlations were validated in an independent set of CRCs with matched methylation and expression data (*n* = 407 COAD and READ samples from The Cancer Genome Atlas, TCGA; Pearson correlation coefficients |r| > 0.5; Figure S11; Table S9; *KLF7* was not analyzed due to missing methylation data).

**Figure 4.**
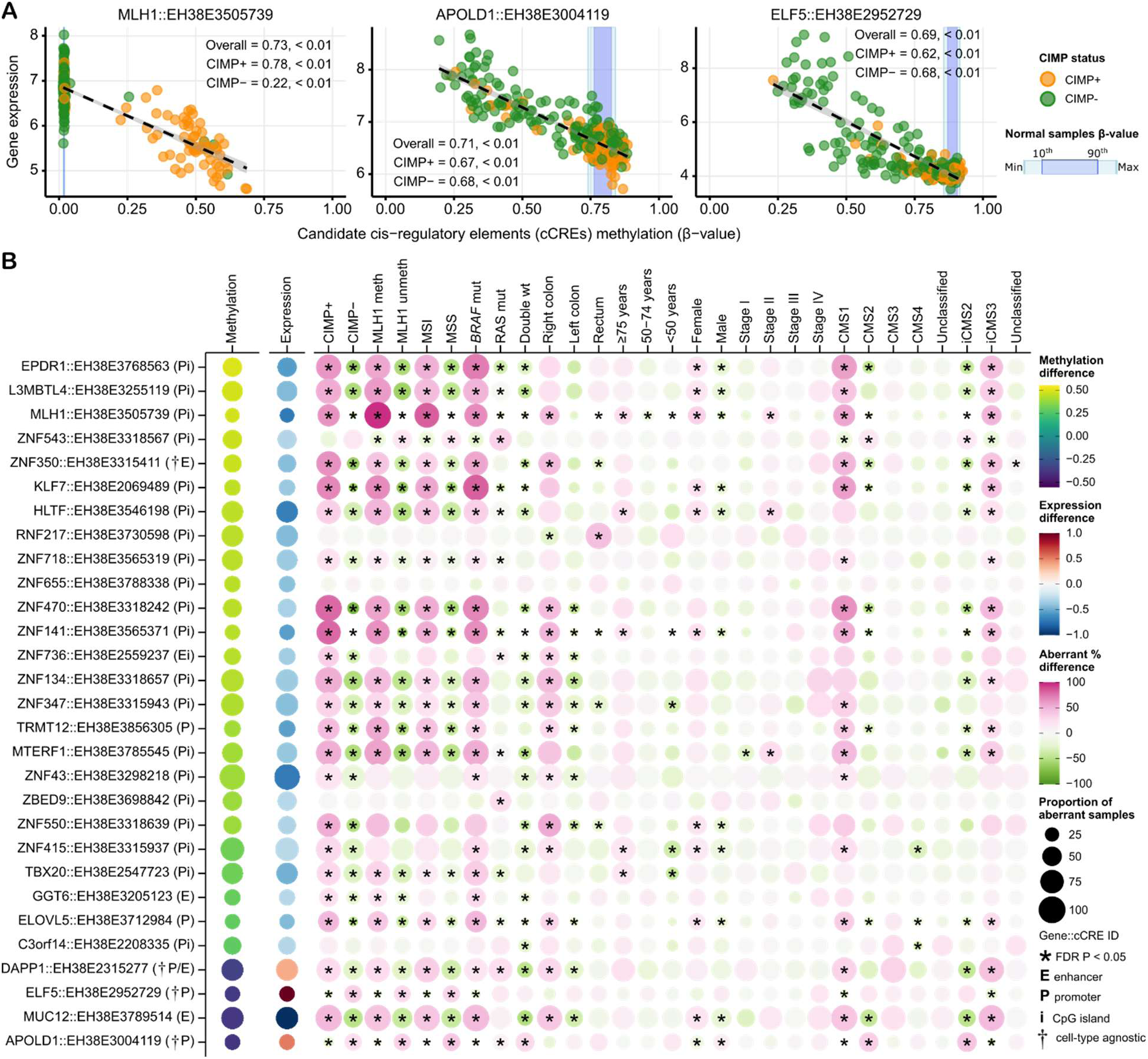
Molecular and clinicopathological associations of intra-tumor heterogeneity (ITH)-low genes with expression regulated by aberrant methylation. **(A)** Scatter plots of representative genes with correlated methylation levels of candidate cis-regulatory elements (cCREs) and target gene expression. Genes were selected to illustrate distinct correlation patterns, including low methylation and high expression in normal and hypermethylation and lower expression in a subset of tumors (MLH1; most common), as well as high methylation and low expression in normal and hypomethylation and higher expression in a subset of tumors (*APOLD1* and *ELF5*). Models were fitted using linear mixed-effects regression, and marginal R^2^ and *P* values are reported for all samples and stratified by CIMP status. Points are colored by CIMP status. Blue shaded regions indicate the methylation range in normal samples (minimum-maximum and 10^th^-90^th^ percentiles). Black dashed lines represent linear model fits with 95% confidence intervals. **(B)** Bubble plot summarizing significant cCRE-gene associations, showing differences in methylation and gene expression between aberrant and non-aberrant samples, as well as associations with other molecular and clinicopathological characteristics. For molecular and clinicopathological variables, bubble size represents the proportion of samples classified as aberrant within each category, and color represents the aberrant percentage-point difference between the selected variable and all other variables within each category. Associations were tested using patient-adjusted logistic mixed-effects models with FDR correction. Asterisks indicate FDR < 0.05. CTA, cell type-agnostic.

As expected, most principal methylation-expression pairs (25 of 29, 86.2%; Figure 4B) showed promoter hypermethylation in a subset of tumors relative to healthy colonic mucosa samples, accompanied by a lower gene expression level. This included the archetypical gene *MLH1* (validated by quantitative methylation-specific PCR) and 13 transcription factor genes, including 12 zinc finger proteins and the Krüppel-like factor *KLF7*. In contrast, four genes, *APOLD1*, *DAPP1*, *ELF5,* and *MUC12*, had hypomethylation of regulatory elements in a subset of tumors compared to healthy mucosa. Hypomethylation was associated with higher expression of *APOLD1*, *DAPP1*, and *ELF5* in the corresponding tumor subsets, whereas *MUC12* inconsistently showed reduced gene expression despite hypomethylation at its predicted enhancer (Figure S9). The regulatory element of *DAPP1* was classified as an active promoter in the cell type-agnostic map but as an enhancer in the large intestine-specific map, and its hypomethylation was unexpectedly enriched in CIMP+ tumors. The regulatory elements of *APOLD1* and *ELF5* were also annotated as cell type-agnostic promoters, but as genomic regions with low chromatin accessibility rather than as active promoters in the large intestine, potentially explaining their aberrant methylation and expression patterns in a subgroup of predominantly CIMP-CRCs. The predicted promoters of both genes lacked CpG islands, a feature consistent with the more tissue-restricted and methylation-variable regulation often observed at CpG-poor promoters.

### Promoter hypomethylation and overexpression of APOLD1 and ELF5 define distinct subsets of canonical CRC

Tumors were classified as aberrant for the methylation-regulated ITH-low genes when the methylation β-values were distinct from the normal mucosa samples (Figure S12; Table S9). There were substantial overlaps among tumors that were aberrant for the different hypermethylated target genes, and nearly all these genes were aberrant predominantly in CIMP+ tumors and/or associated with canonical CIMP+ features in both the in-house cohort and TCGA, including *MLH1* promoter methylation, MSI, *BRAF* V600E mutation, right-sided tumor location, older age, female sex, and the transcriptomic subtypes CMS1 and iCMS3 (Figures 4B and S13; Table S10). This was supported by transcriptomic gene set enrichment analysis, showing that aberrant tumors for all genes had high signals for BRAF-like, MSI-like, immune, and secretory Paneth-like cell signatures (Figure S14; Table S11). In contrast, tumors aberrant for *APOLD1* and *ELF5* had distinct molecular and clinicopathological associations, and promoter hypomethylation was primarily found in CIMP-, MSS, CMS2 and iCMS2 tumors (Figures 4B and S13; Tables S10). Aberrant *APOLD1* was most frequent in *BRAF* and RAS wild-type tumors, while aberrant *ELF5* was associated with RAS hotspot mutations. Both were underrepresented among right-sided tumors. Accordingly, aberrant reactivation of these two genes by promoter hypomethylation are distinct features of subsets of canonical primary CRCs (Figure 5A).

**Figure 5.**
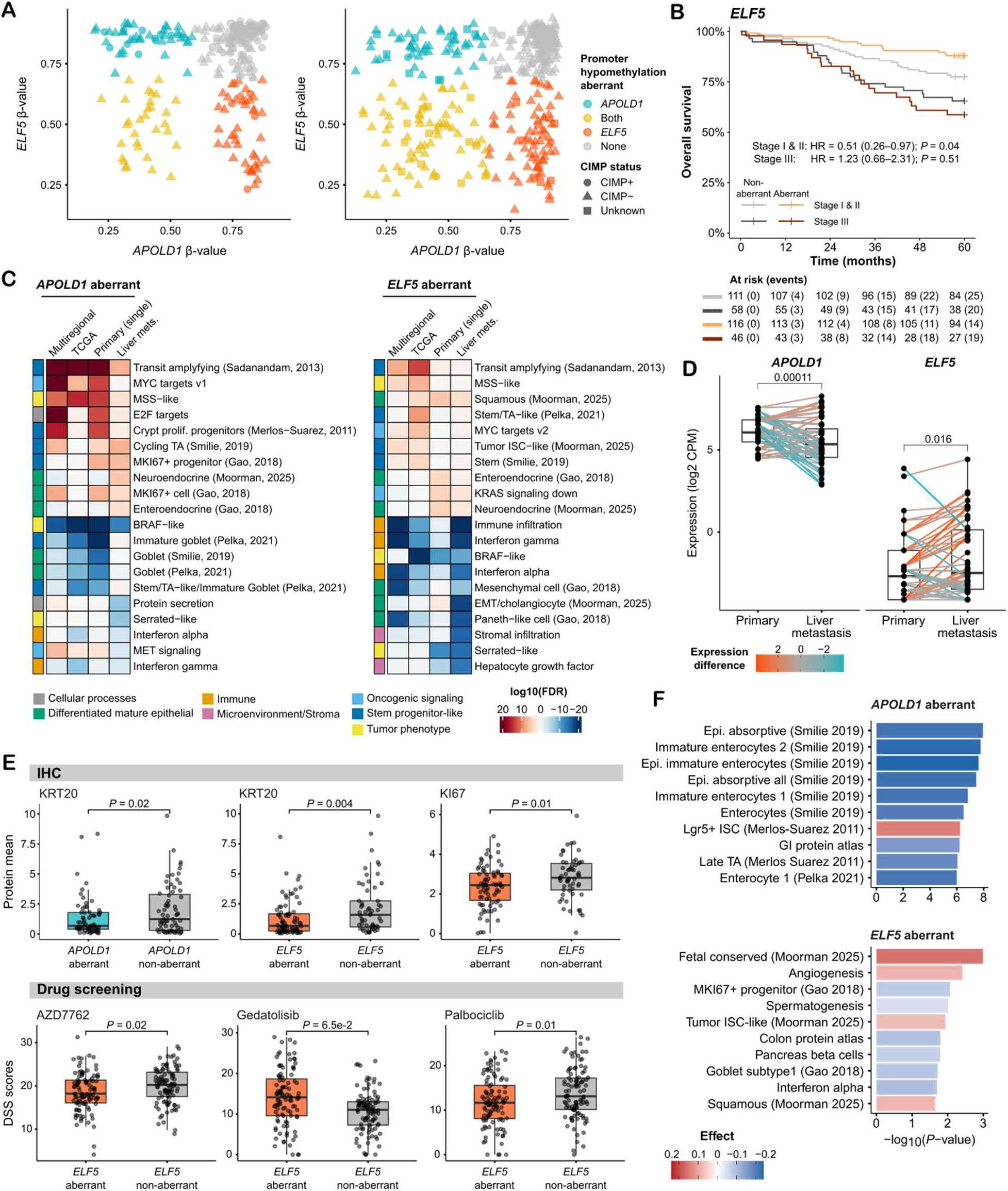
Aberrant *APOLD1* and *ELF5* identify distinct stem-like subgroups of canonical CRC. **(A)** Scatter plots of promoter methylation β-values for *APOLD1* and *ELF5* in the in-house multiregional (left; *n* = 274) and TCGA (right; *n* = 407) CRC cohorts. Points are colored by promoter hypomethylation status of each gene and shapes indicate CIMP status. **(B)** Kaplan-Meier survival curves of overall survival stratified by *ELF5* aberrant status and tumor stage (I-II versus III) in CIMP- and/or MSS CRCs from the multiregional and extended in-house series (*n* = 331). Hazard ratios (HRs), 95% confidence intervals, and *P* values are shown for aberrant versus non-aberrant groups. **(C)** Heatmaps of gene set enrichment analyses for CIMP- and/or MSS samples in each cohort. Colors indicate signed -log10(FDR) values from comparisons of aberrant versus non-aberrant tumors for each gene. The top 10 up- and down-regulated gene sets ranked by mean enrichment significance across the three primary CRC cohorts and from the liver metastasis cohort separately are illustrated. **(D)** Boxplots with paired individual data points of *APOLD1* and *ELF5* expression in patient-matched primary tumors and liver metastases profiled by RNA sequencing (*n* = 21 patients; 81 samples, all CIMP- and/or MSS). Lines connect paired samples and are colored by the expression difference (metastasis versus primary). *P* values were calculated using paired *t* tests. **(E)** Boxplots with overlaid jitter showing selected immunohistochemistry (IHC) protein markers and drug responses in patient-derived organoids (PDOs; *n* = 213) from MSS liver metastases. *P* values were derived from linear models testing associations between each variable and gene aberrant status, adjusting for expression platform (RNA sequencing and microarrays). **(F)** Bar plots of top 10 significantly enriched gene sets from comparisons of PDOs with aberrant versus non-aberrant *APOLD1* and *ELF5* status. Bars represent - log10(*P* value) and are colored by effect size from linear models adjusted for expression platform. In boxplots, center lines indicate medians, boxes represent interquartile ranges (IQRs), and whiskers extend to 1.5x IQR. For cohorts lacking methylation data, gene aberrant status was defined using a cohort-specific median expression cutoff.

Further investigation of molecular and clinical associations was performed in larger and previously published in-house transcriptomic datasets of primary tumors (*n* = 357 CIMP- and/or MSS samples) and liver metastases (*n* = 339 samples). Here, aberrant *APOLD1* and *ELF5* status was defined based on expression above the median in each sample set (Figure S15). Survival analyses of stage I-III CIMP- and/or MSS CRCs treated by complete (R0) surgical resection in the combined multiregional and extended primary tumor samples set (*n* = 331 patients) revealed better overall survival with aberrant *ELF5* in patients with stage I-II CRC, but no prognostic association in stage III (Figure 5B). The prognostic association in early-stage cancers was maintained in multivariable Cox regression models with patient sex, age, and tumor location (multivariable hazard ratio 0.49, *P* value = 0.039; Figure S17). *APOLD1* expression was not associated with survival in any cancer stage (Figure S16 and S17).

There was no pattern of co-occurrence of promoter hypomethylation of *APOLD1* and *ELF5* among tumors in either the in-house cohort or TCGA (Figure 5A), and gene set enrichment analyses among CIMP- and/or MSS tumors demonstrated partially overlapping but distinct biological programs for the two genes (Figure 5C). Both *APOLD1* and *ELF5* aberrant samples were enriched with stem-like cell transcriptomic signatures, particularly with transit amplifying-like cells, although this feature was strongest for *APOLD1.* The main distinctions were that tumors with the *APOLD1* phenotype showed high cell cycle activity and enrichment with proliferative cells, as well as strong depletion of differentiated epithelial signatures, while *ELF5* aberrant tumors were enriched with a trans-differentiation signature of squamous-like cells and were also depleted of immune-related signatures (Figure 5C; Table S12). Notably, these associations were similar among primary tumors and liver metastases, although the expression levels of the two genes varied during metastatic progression. *ELF5* was generally expressed higher in liver metastases, but comparisons of matched primary-metastasis samples showed substantial variation among patients. Some patients with aberrant *ELF5* in the primary tumor showed low or moderate expression levels in their metastases, and vice versa (Figures 5D and S18).

Functional consequences were further explored in patient derived organoids (PDOs) of liver metastases with available transcriptomic (*n* = 206), drug response (*n* = 24 drugs in 204 PDOs), and fluorescence-based immunohistochemistry data (*n* = 14 proteins in 135 PDOs; Figure 5E-F; Table S13). Gene set enrichment analyses supported the stem-like features of both *APOLD1* and *ELF5* aberrant tumors, with depletion of differentiated cell types and enrichment with intestinal stem-like cells in both groups of PDOs (Figure 5F). Immunohistochemistry further confirmed lower protein expression levels of the differentiation marker KRT20 in both groups, as well as of CDX2 in *APOLD1* aberrant PDOs. Aberrant *ELF5* PDOs were also depleted of transcriptomic signals for proliferative MKI67+ progenitor cells and showed lower expression levels of the KI67 proliferation marker at the protein level. Consistently, these PDOs were also less sensitive to drugs targeting cell cycle progression, including the checkpoint kinases (Chks) inhibitor AZD7762 and the CDK4/6 inhibitor palbociclib, supporting that aberrant *ELF5* is a marker of a slow-cycling, stem-like state. The transcriptomic analyses further indicated strong enrichment with fetal-like programs in *ELF5* aberrant PDOs, along with the squamous cell-like trans-differentiation signature, suggesting a role of aberrant *ELF5* also in phenotypic plasticity. The strongest pharmacological association was sensitivity to the dual PI3K/mTOR inhibitor gedatolisib in the *ELF5* aberrant group (Figure 5E; Table S13), potentially reflecting the role of mTOR signaling in cellular quiescence and the sensitivity of cancer stem cells to mTOR inhibition^27^.

## Discussion

This study identified promoter hypomethylation linked to aberrant cell states in MSS CRC. Focal hypomethylation events and/or overexpression of genes that are normally silent or expressed at low levels in the colonic mucosa were associated with transcriptomic signatures of cellular de-differentiation and trans-differentiation in both primary and metastatic tumors, and with supporting protein and drug sensitivity data in tumor organoids. This showed that methylation-expression associations in CRC extend beyond the well-established link between promoter hypermethylation and gene silencing in CIMP+ tumors. Specifically, unconventional hypomethylation events occurred in a substantial proportion of otherwise canonical CRCs with conventional molecular features, including CIMP-, MSS, CMS2, iCMS2, and *BRAF* wild-type status. These findings suggest a two-level model of epigenetic organization in CRC. At one level, CIMP represents a genomically broad and spatially ubiquitous tumor phenotype established early in tumor development and aligned with MSI-like biology. In parallel, focal methylation changes, particularly promoter hypomethylation, are associated with transcriptomic cell state transitions in tumors that otherwise share canonical molecular characteristics. Such focal events remain capable of change during disease progression and might be involved in phenotypic plasticity, as suggested by expression discordances in patient-matched samples of primary and metastatic tumors.

We identified two target genes of unconventional hypomethylation events. Transcriptomic activation of *APOLD1* and *ELF5* occurred in distinct but partially overlapping subgroups that combined accounted for more than half of CIMP-tumors (approximately 20% each, plus 17% overlap). Both subgroups showed de-differentiation into stem-like and transit amplifying-like states, the latter representing an early intermediate cell state in tissue regeneration^28^. *APOLD1* aberrant tumors further showed a proliferative phenotype. *APOLD1* encodes an endothelial early-response protein that is involved in vascular homeostasis and hypoxia response^29^. However, aberrant APOLD1 expression was not accompanied by increased VEGF-pathway or angiogenic activity in our study, consistent with a previous study of MSS CRC^30^. Transcriptomic associations were also retained in PDOs, which lack a cellular vasculature, suggesting that aberrant *APOLD1* expression reflects a proliferative stress response of malignant epithelial cells in CRC. In contrast, *ELF5* aberrant tumors showed a slow-cycling phenotype accompanied by fetal de-differentiation and transition to a non-canonical intestinal state. Consistently, *ELF5* encodes an epithelial transcription factor involved in cell-fate specification, and has recently been linked to a squamous-like trans-differentiation program that emerges in CRC as the cancer cells lose canonical intestinal identity^3^. This relatively slow-cycling phenotype is also consistent with the favorable prognostic association of *ELF5* in early-stage disease, although its expression level increased during metastasis, suggesting that *ELF5*-associated states may be remodeled and have different roles during the stages of metastatic progression. Notably, we did not establish a functional role of the gene products in governing cell state transitions, and this study can only propose these genes as markers of cellular plasticity.

Promoter hypomethylation associated with gene expression activation is not unique to CRC, but despite longstanding interest in this mechanism, relatively few examples have been identified, and their functional and clinical relevance remains less well established than for hypermethylation-associated gene silencing. Specific examples include hypomethylation-mediated activation of *PAX2* in endometrial carcinogenesis^31^, elevated *HIF1A* expression in aggressive breast cancer phenotypes^32^, and activation of the long non-coding RNA *HUMT* promoting lymphangiogenesis and metastasis in triple-negative breast cancer^33^. Genome-wide low promoter methylation has been associated with oncogenic subtypes and adverse outcomes in T-cell acute lymphoblastic leukemia^34^. However, few studies have implicated recurrent focal promoter hypomethylation events in cell state transitions, and this study extends the concept of cancer-associated hypomethylation towards a potential role in phenotypic diversification of canonical CRCs.

Several aspects of the analytical approach were important for detecting robust cancer cell-intrinsic methylation-expression relationships in this study. First, a broader regulatory annotation of methylation probes on the EPIC array substantially expanded the search space outside conventional gene-centric promoter annotations to include regulatory elements with potential tissue- and cell-state specific activity^21,35^. This was particularly relevant in the context of tumor plasticity analysis, where changes in transcriptomic states may involve regulatory elements outside canonical CpG-island promoters. The extended annotation also improved the predictions of target genes for each regulatory element. Second, the multiregional tumor sampling approach enabled distinction between cancer cell-associated methylation-expression relationships from signals driven mainly by intratumoral variation of the microenvironment. This is particularly important in bulk tumor analyses, where varying proportions of immune, stromal, endothelial, and malignant cells can generate methylation-expression associations that reflect differences in cell type compositions rather than regulatory mechanisms within cancer cells. Reproducibility of key expression phenotypes in PDO cultures of malignant epithelial cells strengthened the confidence that these features reflect cancer cell-intrinsic biology.

The main limitation of the study is the associative design and lack of functional perturbation experiments to validate epigenetic regulatory mechanisms and their role in controlling cell state transitions. However, the conclusions of this study were supported by validation of methylation-expression regulatory associations in an independent set of tumors, and by orthogonal functional approaches in PDOs. Immunohistochemistry of differentiation and proliferation markers, as well as drug screen data for cell cycle and stem cell inhibitors supported the dedifferentiated and proliferative transcriptomic phenotypes. Particularly, mTOR inhibition is a proposed strategy to target cancer stem cells, consistent with the strong sensitivity in *ELF5* aberrant PDOs^27^. Furthermore, although our approach to deconvolute cancer cell-intrinsic signals by multiregional tumor sampling allows scalability to large tumor series, single-cell methylome-transcriptome profiling will be needed to resolve directly the relationship between regulatory methylation states and transcriptomic phenotypes within individual malignant cell populations. The feasibility of this approach in CRC has been demonstrated in a previous joint single-cell genome, methylome, and transcriptome study showing methylation heterogeneity across lineages and linking distinct epigenetic states to tumor evolution^36^. However, single-cell DNA methylation profiling has not yet reached the methodological maturity or routine scale of single-cell RNA sequencing. Unlike gene-centered transcriptomic count matrices, single-cell DNA methylation sequencing produces sparsely sampled global methylation data, often requiring aggregation across regions and specialized analytical approaches to distinguish biological heterogeneity from differences in coverage^37^. Current protocols also remain labor-intensive^38^. Thus, multiregional bulk tumor sampling combined with reannotation of regulatory elements can provide a complementary strategy to identify reproducible methylation-expression relationships in translational studies, while future single-cell methylome studies will be required to resolve the cellular heterogeneity underlying these associations. We also expect that investigating epigenetic data layers beyond DNA methylation will uncover additional regulatory mechanisms of phenotypic plasticity.

In conclusion, this study shows that focal promoter hypomethylation is associated with transcriptomic phenotypes distinct from the established phenotype of broad promoter hypermethylation in CRC. These focal events can pinpoint subgroups of canonical CRCs with distinct cellular plasticity programs.

## Resources availability

### Lead contact

Requests for further information and resources should be directed to the lead contact, Anita Sveen.

### Materials availability

This study did not generate new unique reagents.

### Data and code availability

Raw and processed EPIC array methylation data generated in this study for the multiregional primary CRC cohort (n = 274) have been deposited in the NCBI’s Gene Expression Omnibus (GEO) under accession number GSE353142. The final probe annotations with the regulatory annotations derived from ENCODE v4 cCREs are provided in Table S2. Transcriptomics data from the same samples have previously been deposited to GEO under accession number GSE241101. Publicly available methylation and gene expression data from TCGA were retrieved using the *TCGAbiolinks* v2.38.0 R package. Transcriptomics data from the extended single-sample primary tumor cohort were previously deposited to GEO under accession numbers GSE24550, GSE29638, GSE69182, GSE79959, GSE96528, GSE139170 and GSE178120, data from the liver metastases cohort under accession number GSE159216, and data from PDOs under accession number GSE2945511 and in Mendeley Data under DOI 10.17632/hr94h42xdc.3. This paper does not report original computer code, and all software and algorithms are described and cited in the manuscript text.

## Acknowledgments

The study was funded by grants from the Norwegian Cancer Society (project numbers 249654 to A.S. and 223319 to R.A.L.), the South-Eastern Norway Regional Health Authority (project numbers 2023101 to A.S. and 2024108 to R.A.L.).

## Author contributions

Conceptualization, L.N., G.E.L., R.A.L., A.S.; methodology, L.N., R.P., G.E.L., R.A.L., A.S.; Investigation, L.N., I.A.E., H.M.V., K.K., A.S.; writing – original draft, L.N. and A.S.; writing – review & editing, all authors; funding acquisition, A.S. and R.A.L..; resources, U.S., O.S.J, S.Y., A.N., K.K., R.A.L.; supervision, A.S. All authors have read and approved the manuscript ahead of submission.

## Declaration of generative AI and AI-assisted technologies in the manuscript preparation process

During the preparation of this work, the authors used GPT UiO to improve the clarity and quality of the text. After using this tool, the authors reviewed and edited the content as needed and take full responsibility for the content of the publication.

## Supplemental information

Document S1. Supplemental Figures S1-S18

Document S2. Supplemental Tables S1-S13

## Methods

### Patient material

This study was approved by the Regional Committee for Medical and Health Research Ethics, South Eastern Norway (REC numbers 1.2005.1629, 2010/1805, and 2017/780). Participants provided written informed consent, and the research was conducted in alignment with the Declaration of Helsinki. Patients were included from an ongoing observational study of CRCs treated by major surgical resection of the primary tumor at Oslo University Hospital. Patients in the current study (*n* = 93 patients and 95 tumors, two patients had synchronous tumors) were treated between 2015-2016, and 2-4 multiregional samples were collected from spatially distinct areas of each tumor (*n* = 276 samples). Healthy colonic mucosa samples were collected from the surgical specimen of each patient, and a set of samples were included for DNA methylation profiling (*n* = 19). Furthermore, one patient (*n* = 2 samples) was excluded due to poor quality of the DNA methylation data (abnormal β-value distribution; Figure S1). The final multiregional tumor sample set included 274 samples from 94 tumors and 92 patients (mean of 2.9 samples per tumor).

### Previously published in-house data

DNA and RNA from the multiregional samples have previously been extracted using Qiagen’s AllPrep DNA/RNA/miRNA Universal Kit (Qiagen GmbH, Hilden, Germany) following the manufacturer’s instructions^18^. The samples have also been analyzed for MSI status using the Promega MSI Analysis System (Promega, Madison, WI, USA), transcriptomic profiling by Affymetrix Human Transcriptome 2.0 arrays (Thermo Fisher Scientific, Waltham, MA, USA) and mutation profiling of *KRAS* and *NRAS* exons 2-4, as well as *BRAF* V600E hotspots by Sanger sequencing (Applied Biosystems, Waltham, MA, USA)^18^. Tumor sample classifications according to CMS, CRIS, iCMS were extracted from the publication^18^. Transcriptomic raw intensity CEL-files (Gene Expression Omnibus accession number GSE241101) were processed as previously described^18^ but with updated CDF file from Brainarray (hta20hsgencodegcdf_25.0.0) and gene annotation from GENCODE (gencode.v36.annotation.gtf).

Additional in-house and previously published transcriptomics data from CIMP- or microsatellite stable (MSS) primary colorectal tumors (*n* = 357 samples; Gene Expression Omnibus accession numbers GSE24550, GSE29638, GSE69182, GSE79959, GSE96528, GSE139170 and GSE178120) and colorectal liver metastases from patients treated by hepatic resection (*n* = 339 samples from 197 patients; GSE159216) were included for analysis^39,40^. Samples were analyzed with Affymetrix GeneChip Human Exon 1.0 ST arrays (*n* = 172 primary tumors) or Human Transcriptome 2.0 arrays (*n* = 185 primary tumors and all 339 liver metastasis samples). Transcriptomic raw intensity CEL-files were processed as previously described^18^, separately for each array platform among primary tumors and metastases group, but with updated CDF file from Brainarray (hue×10sthsgencodegcdf_25.0.0 and hta20hsgencodegcdf_25.0.0) and gene annotation from GENCODE (gencode.v36.annotation.gtf). For the primary tumors, only genes represented on both array platforms were retained before the datasets were combined and batch-corrected using the *ComBat* function from the *sva* v3.60.0 R package^41^. Patient-matched primary tumor and metastasis sample sets were available for 44 patients (total *n* = 23 primary and 79 metastasis samples analyzed on Human Transcriptome 2.0 arrays and *n* = 21 primary and 60 metastasis samples analyzed with RNA sequencing), with mean expression values used for primary tumors with multiregional sampling.

### CpG-island methylator phenotype (CIMP) determination

*MLH1* promoter methylation and CIMP status based on the Weisenberger marker panel (*CACNA1G*, *RUNX3*, *IGF2*, *SOCS1*, *NEUROG1*) were analyzed using quantitative methylation-specific PCR with the ALU-C4 repetitive element for reference, as previously described^19^. For each sample, 900 ng of genomic DNA was bisulfite converted using the EpiTect Bisulfite Kit (Qiagen) according to the manufacturer’s standard protocol. The converted DNA was automatically purified on a QIAcube instrument (Qiagen) and eluted in a final volume of 40 μL (2 × 20 μL). Primers and probes were from BioNordika (Oslo, Norway) and Thermo Fisher Scientific, respectively. Methylation levels were estimated as the percentage of methylated reference (PMR) and genes with PMR ≥10 were considered methylated. Samples with three or more methylated marker genes were classified as CIMP-positive (CIMP+)^10^. CIMP data have previously been published for a subset of the tumors (*n* = 30)^19^.

### DNA methylome profiling

For each sample, 500 ng of genomic DNA was subjected to bisulfite conversion using the EZ-96 DNA Methylation Kit (Zymo Research, Irvine, CA, USA) following the manufacturer’s protocol. The samples were further processed according to the Infinium HD Assay Methylation Protocol Guide, hybridized onto an Infinium MethylationEPIC BeadChip and scanned using the iScan system (Illumina, San Diego, CA, USA). Initial quality control of the intensity data files was performed using the GenomeStudio v2011.1 with the Methylation Module v1.9 supplemented by the BeadArray Controls Reporter software v1.1.0 (Illumina). Raw data processing and analysis were performed as previously described^42^, using *R* v4.5.2 and packages from *Bioconductor* (release 3.22). Raw intensity data (IDAT) files were preprocessed using the *minfi* v1.56.0 package^43^ with the *preprocessFunnorm* function for data normalization^44^. Probes outside the detection threshold (*P* value > 0.01), located on sex chromosomes, containing a single nucleotide polymorphism at the CpG interrogation site or at the single nucleotide extension site (minor allele frequency, MAF ≥ 0.01), and deemed cross-reactive (as identified by the *maxprobes/dropXreactiveLoci* v0.0.2 list^45,46^) were excluded from further analysis. β-values (proportion of methylated DNA at each CpG site) and corresponding M-values (logit-transformed β-values) were extracted for the retained probes. Batch effects were analyzed with Singular Value Decomposition (SVD) as implemented in *ChAMP* v2.40.0 Bioconductor package^47^. However, it was not possible to distinguish biological variation from potential technical batch effects due to unbalanced distribution of samples across arrays, with adjacent localization of multiregional samples from the same patients and collection of all normal samples on the same arrays. Batch correction was therefore not performed to avoid the risk of removing true biological signals and introduce bias.

### Re-annotation of EPIC array methylation probes

Probe annotations for the Illumina EPIC array were updated to the GRCh38/hg38 human reference genome using annotation data from the *InfiniumAnnotation* resource (https://zwdzwd.github.io/InfiniumAnnotation). Candidate cis-regulatory elements (cCREs) from ENCODE v4 were downloaded from the SCREEN web interface (https://screen-v4.wenglab.org) in February 2025^22^. We retrieved both the general and cell type-agnostic classification of all human cCREs, as well as the biosample-specific cCREs for the large intestine (*n* = 7 primary tissue samples and 2 cell lines in the Core Collection containing assay data for DNase hypersensitive sites, H3K4me3 or H3K7ac marks, and binding of CTCF). Cell type-agnostic and large intestine-specific cCREs were mapped to the methylation probes based on genomic coordinates.

For the large intestine-specific annotation, each cCRE was assigned a regulatory classification based on the most frequently observed category across the nine ENCODE tissues or cell lines. The category “Low-DNase” was excluded if any alternative classifications were available. In the event of a tie between remaining categories, a predefined priority hierarchy was applied in the following order: promoter-like signature (PLS), proximal enhancer-like signature (pELS), distal enhancer-like signature (dELS), candidate poised promoter (Ca-H3K4me3), CTCF-bound element (CA-CTCF), transcription factor-bound element (CA-TF), and candidate element with no other marker (CA-only).

Predicted target genes for each cCRE were retrieved from the SCREEN database. These predictions were based on the four computational models ABC^23^, EPIraction^24^, GraphRegLR^25^, and rE2G^26^. Repetitive genomic elements were annotated separately by mapping UCSC RepeatMasker data (rmsk_hg38_USCS.txt.gz) to the probe coordinates.

Final annotations were assigned using three complementary frameworks. The Illumina default annotation classified probes into promoters (including TSS1500, TSS200, 1^st^ exon, or 5’ UTR based on the *UCSC_RefGene* or *GencodeCompV12* columns), enhancers (located within *Phantom4*, *Phantom5* or *x450K* regions), dual (overlapping both promoter and enhancer regions), gene bodies (Body or 3’ UTR), and intergenic regions (any remaining probes). The ENCODE cell type-agnostic annotation, in which probes were categorized based on ENCODE-defined cCREs, included promoters (PLS), enhancers (pELS or dELS), poised promoters (CA-H3K4me3), CTCF-bound elements (CA-CTCF), transcription factor-bound element (CA-TF), repeats (LINE, SINE, LTR or DNA repeats), gene bodies (based on distance to transcription start site), and intergenic elements. The ENCODE large intestine-specific annotation was based on the same criteria as the cell type-agnostic annotation but was restricted to large intestine-specific cCREs, allowing for analysis of the tissue-specific regulatory context.

### Intra-tumor transcriptomic and methylation heterogeneity

Transcriptomic and DNA methylation-based intra-tumor heterogeneity (ITH) among multiregional tumor samples were quantified separately, following a previously described approach^18^. Initially, probe-wise differential methylation analysis was performed between tumor and normal samples using the *lmfit* and *eBayes* functions from *limma* v3.66.0^48^ on the methylation M-value matrix. Delta β (Δβ) values were computed using the R package *ABC.RAP* v0.9.0 based on a groupwise comparison of β-values between tumor and normal samples. Probes with Δβ > 0.1 and Δβ < -0.1 (Benjamini-Hochberg adjusted *P* < 0.05) were classified as hypermethylated and hypomethylated, respectively, and kept for downstream analyses. Genes with low expression variation (10-90^th^ percentile range < 1) and probes with low methylation variation (10-90^th^ percentile range < 2) across the complete tumor sample set were considered non-informative and excluded. For each retained gene in the expression dataset and probe in the methylation dataset, a linear mixed-effects model was fitted across all samples from all tumors using *lmer* (*lme4* v1.1-38), with “tumor” as a random effect, as previously described^49^. Intra-class correlation coefficients (ICCs) were calculated using the *icc* function in the R package *performance* v0.16.0^50^, and ITH-scores were defined as 1 – ICC. ITH-scores were grouped as ITH-low (≤ 0.3), ITH-intermediate (> 0.3 - < 0.6), and ITH-high (≥ 0.6).

### Transcriptomic and methylation data correlation analysis

Associations between methylation of regulatory elements and gene expression were assessed using probes and genes classified as ITH-low. For DNA methylation data, ITH-low probes located in gene bodies or intergenic regions according to the ENCODE cell type-agnostic and large intestine-specific annotations were excluded. Median methylation values were computed for all probes targeting each regulatory region, and only regions with at least three mapped probes were retained. Each regulatory element was paired with its predicted target gene, provided that the gene was also ITH-low. Pearson’s correlations were estimated and linear mixed-effects models relating gene expression to the methylation level of the corresponding regulatory element were fitted with *lmer* (*lmerTest* v3.2-0)^51^ and using “tumor” as a random effect. Marginal and conditional R^2^ values were calculated using the *r2_nakagawa* function in the R package *performance* v0.16.0^50^, and the regression coefficients and *P* values were extracted from model summaries. Multiple testing correction was applied using the false discovery rate (FDR). Subgroup analyses according to CIMP status were performed with the same approach. Significant correlations were defined as adjusted *P* < 0.01 and marginal R^2^ > 0.5 in at least one of the analyses (overall and by subgroup). To account for repeated associations involving the same cCREs or genes, methylation-expression correlations were ranked by the marginal R^2^. Correlations that were simultaneously ranked as the strongest for both the cCRE and the gene were classified as the main correlations, while all remaining were classified as secondary correlations. For the significant main correlations, tumor samples were identified as aberrantly methylated based on absolute Z-scores > 3 and the absolute Δβ value relative to the mean of normal samples above 0.15. Absolute Z-scores were calculated as:

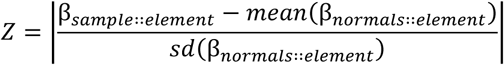

where β represents the β-value for the respective regulatory element.

### TCGA validation cohort data collection

CRC samples from TCGA (COAD and READ projects) with matched DNA methylation and gene expression data were retrieved using the *TCGAbiolinks* v2.38.0 R package^52^. DNA methylation data from the Illumina HumanMethylation450 BeadChip platform were obtained as β-values, while RNA sequencing data were downloaded as gene-level counts generated using the STAR workflow (“STAR – Counts”). Corresponding sample- and feature-level annotations were extracted for analysis. Additional molecular and clinical annotations, including MSI status, CIMP status, *MLH1* promoter methylation and tumor location, were obtained using the *UCSCXenaTools* v1.7.0 R package^53^.

### Functional analyses in tumor organoids

Tumor organoids (*n* = 213) have previously been established from the same observational study as the CRC liver metastases of patients treated by hepatic resection (*n* = 102 patients, including 2-6 metastatic deposits from 65 of the patients)^54^. All organoids are MSS and subsets were analyzed by transcriptomic profiling (*n* = 206 PDOs of which 119 by Affymetrix Human Transcriptome 2.0 arrays, GSE2945511, and 87 by RNA sequencing, Mendeley Data 10.17632/hr94h42xdc.3), drug sensitivity testing (*n* = 24 drugs in 204 PDOs), and multiplex fluorescence-based immunohistochemistry (*n* = 14 proteins in 135 PDOs). Pre-processed data were retrieved from the original study^54^. Immunohistochemistry data were retrieved as the normalized mean relative expression of each protein. Drug sensitivity data were retrieved as drug sensitivity scores (DSS).

### Statistical analysis

All statistical analyses were performed using R v4.5.2^55^. Tumor purity was estimated from gene expression data using the *estimate* v1.0.13 R package^56^. PCAs were performed using the *prcomp* function in R, with features mean-centered and scaled to unit variance. Associations between aberrant DNA methylation status and clinical or molecular characteristics were evaluated using logistic regression, with patient as a random intercept for the multiregional samples, with *P* values adjusted by FDR. Single-sample gene set enrichment analyses of a custom collection (*n* = 157 gene sets; Table S7) were performed using the *GSVA* v2.4.1 package^57^, and differential pathway activity was assessed using the *subCamera* function from the *CMScaller* v2.0.1 package^58^. For the PDOs, group comparisons of molecular data were performed with linear regression models, using aberrant gene status as the primary predictor and analysis platform as a covariate. Survival analyses were performed for patients with stage I-III CRC (including both patients with multiregional and single tumor samples) treated by complete tumor resection (residual tumor status R0) and with no pre-surgical chemoradiation or synchronous tumors. For multiregional samples, a gene was classified as aberrant at the patient level if detected as aberrant in at least one sample. Kaplan-Meier estimates for overall survival, defined as the time from surgery to death from any cause, were generated using the package *survival* v3.8-6. Patients were censored at 5-year follow-up. Cox proportional hazard models were fitted separately by stage (stages I-II versus III) and reported as hazard ratios with 95% confidence intervals and *P* values. Data visualization was performed primarily using *ggplot2* v4.0.2. Specialized plots were generated using the packages *ComplexHeatmap* v2.26.0, *ggridges* v0.5.7, *scattermore* v1.2, *ggalluvial* v0.12.6, *GGally* v2.4.0, *PupillometryR* v0.0.7, *ggsurvfit* v1.2.0, and *survminer* v0.5.2. All statistical tests were performed with two-sided *P* value or FDR < 0.05 as a threshold of significance.

