## Supplemental Figures S1-S18 for "Epigenetic reprogramming by promoter hypomethylation is linked to aberrant gene expression and transcriptomic plasticity in colorectal cancer"

**Supplemental table titles**

**Table S1.** Clinicopathological characteristics of multiregional colorectal tumors across CIMP groups.

**Table S2.** Integrated annotation of EPIC methylation probes with ENCODE regulatory features and predicted target genes.

**Table S3.** Genome-wide differential methylation between tumor and normal samples.

**Table S4.** Differential methylation enrichment statistics across genomic annotations.

**Table S5.** Intra-tumour heterogeneity landscape derived from transcriptomic data.

**Table S6.** Intra-tumour heterogeneity landscape derived from DNA methylation data.

**Table S7.** Associations between principal components and gene set activity by intra-tumor heterogeneity status.

**Table S8.** Methylation-expression correlations across the multiregional cohort.

**Table S9.** Differential methylation and gene expression by aberrant status across the multiregional and TCGA cohorts.

**Table S10.** Clinicopathological associations of gene aberrant status across cohorts.

**Table S11.** Gene set enrichment associations of gene aberrant status across cohorts.

**Table S12.** *APOLD1* and *ELF5* pathway enrichment analyses in CIMP- and/or MSS samples.

**Table S13.** *APOLD1* and *ELF5* associations with molecular and functional in patient-derived organoids.

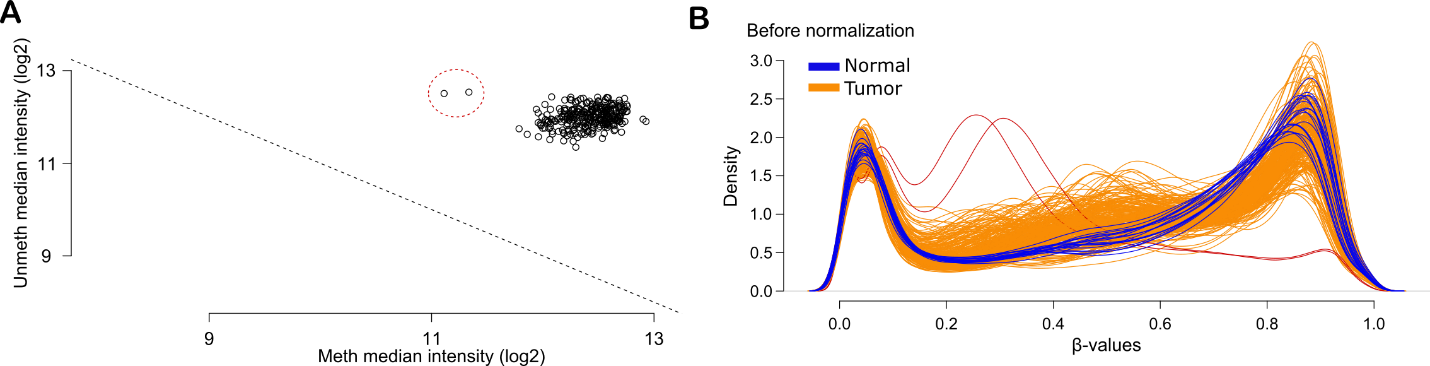
**Supplemental figures**

**Figure S1. Quality control of Illumina EPIC array methylation data. (A)** Scatter plot of log2-transformed median methylated and unmethylated signal intensities across samples. Each point represents one sample. Two outlier samples excluded from further analyses, highlighted by a red dashed circle, cluster separately from the main distribution. **(B)** Density plot of β-values for tumor (orange) and normal (blue) samples before normalization. The two excluded samples (red) showed skewed distribution compared to the remaining samples.

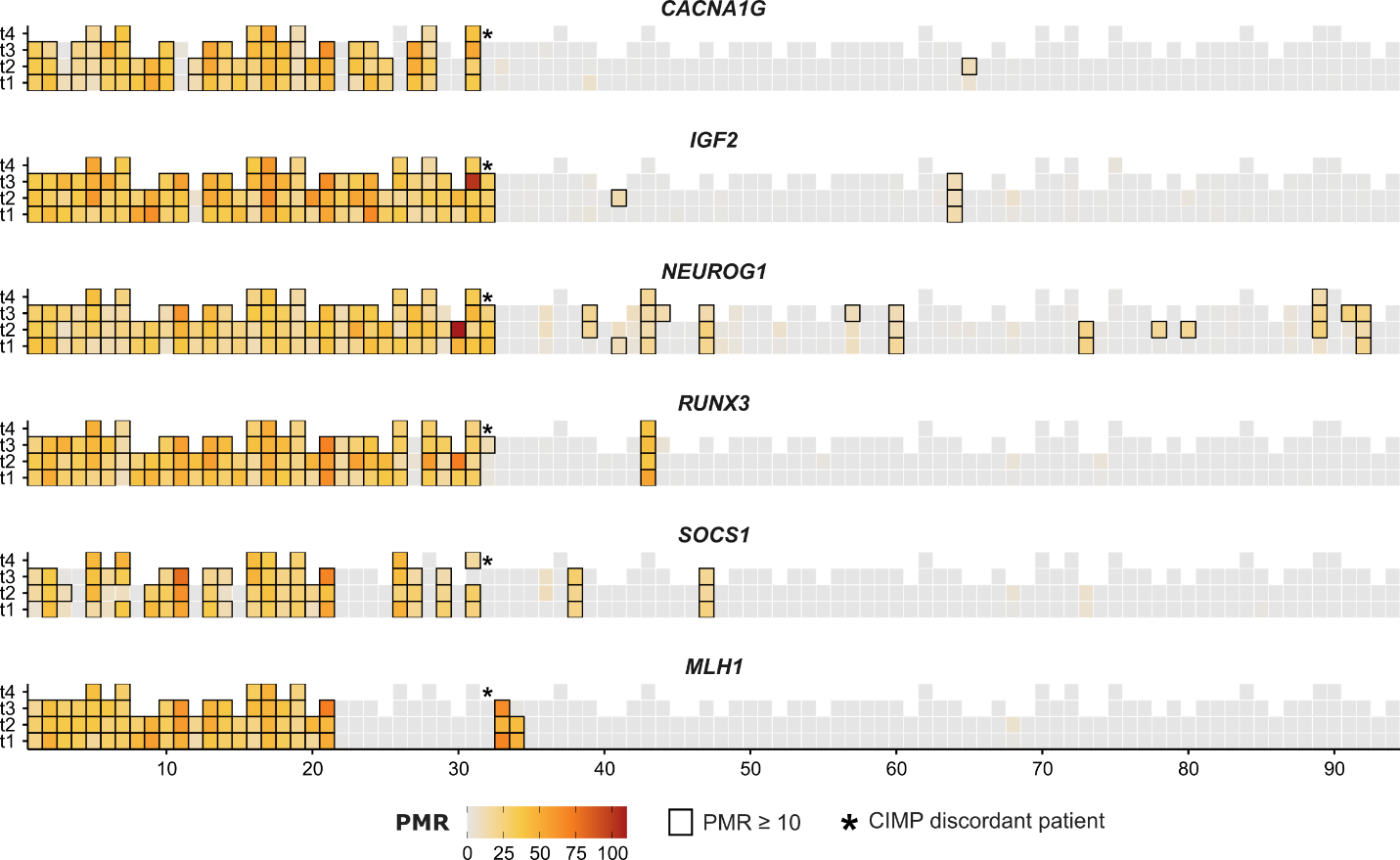
**Figure S2. Methylation levels of the five-marker Weisenberger CIMP panel and *MLH1*.** Each column represents an individual tumor and regions (t1–t4) are color-coded according to their percent methylated reference (PMR) values. Tiles with PMR ≥ 10 are highlighted by a black border.

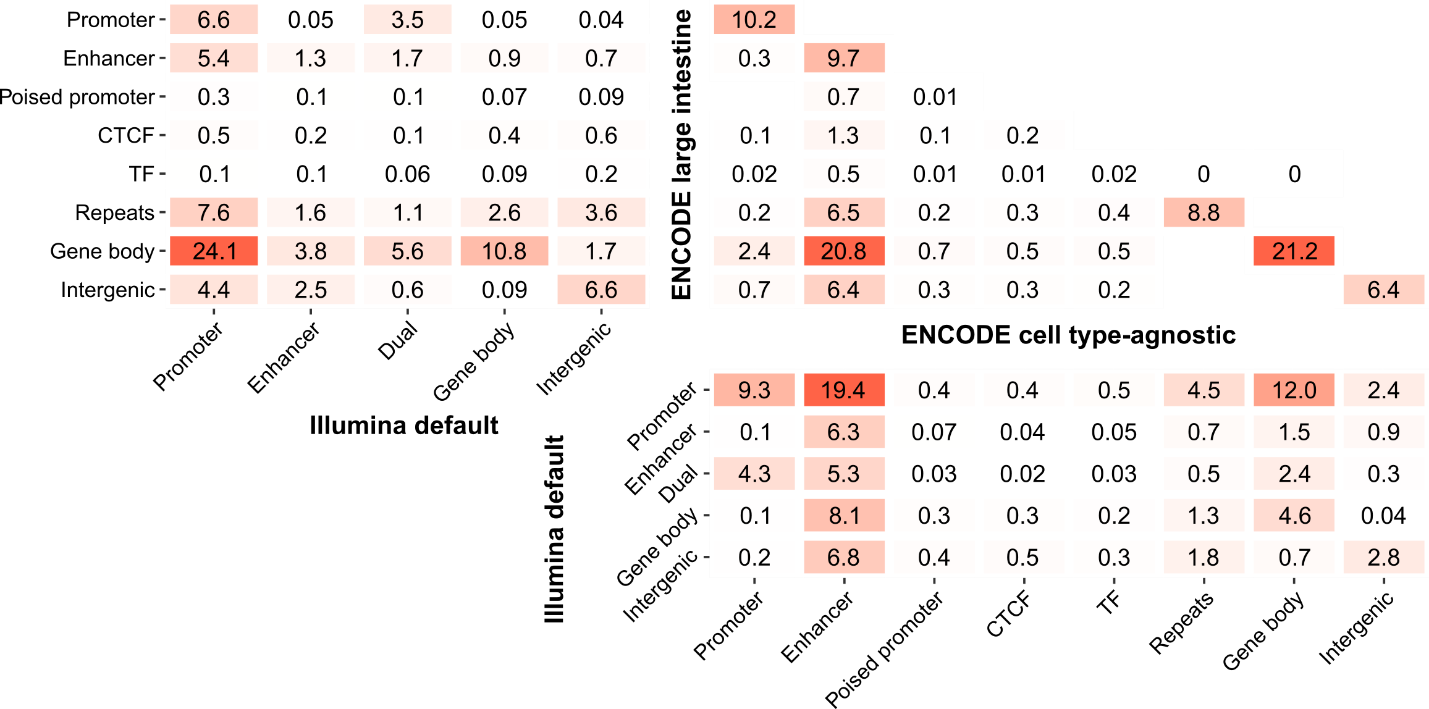
**Figure S3. Comparison of CpG probe annotations between Illumina and ENCODE-derived datasets.** Comparison of CpG probe annotations based on the default Illumina classification and ENCODE-derived annotations, including cell type-agnostic and large intestine-specific categories. Values within boxes indicate the percentage of probes assigned to each category, with color intensity scaled according to the percentage. In the Illumina classification, “dual” denotes probes overlapping both promoter and enhancer regions. In the ENCODE-derived classifications, CTCF denotes CCCTC-binding factor-associated regions, which typically correspond to insulator elements involved in chromatin organization; TF denotes transcription factor-associated regions.

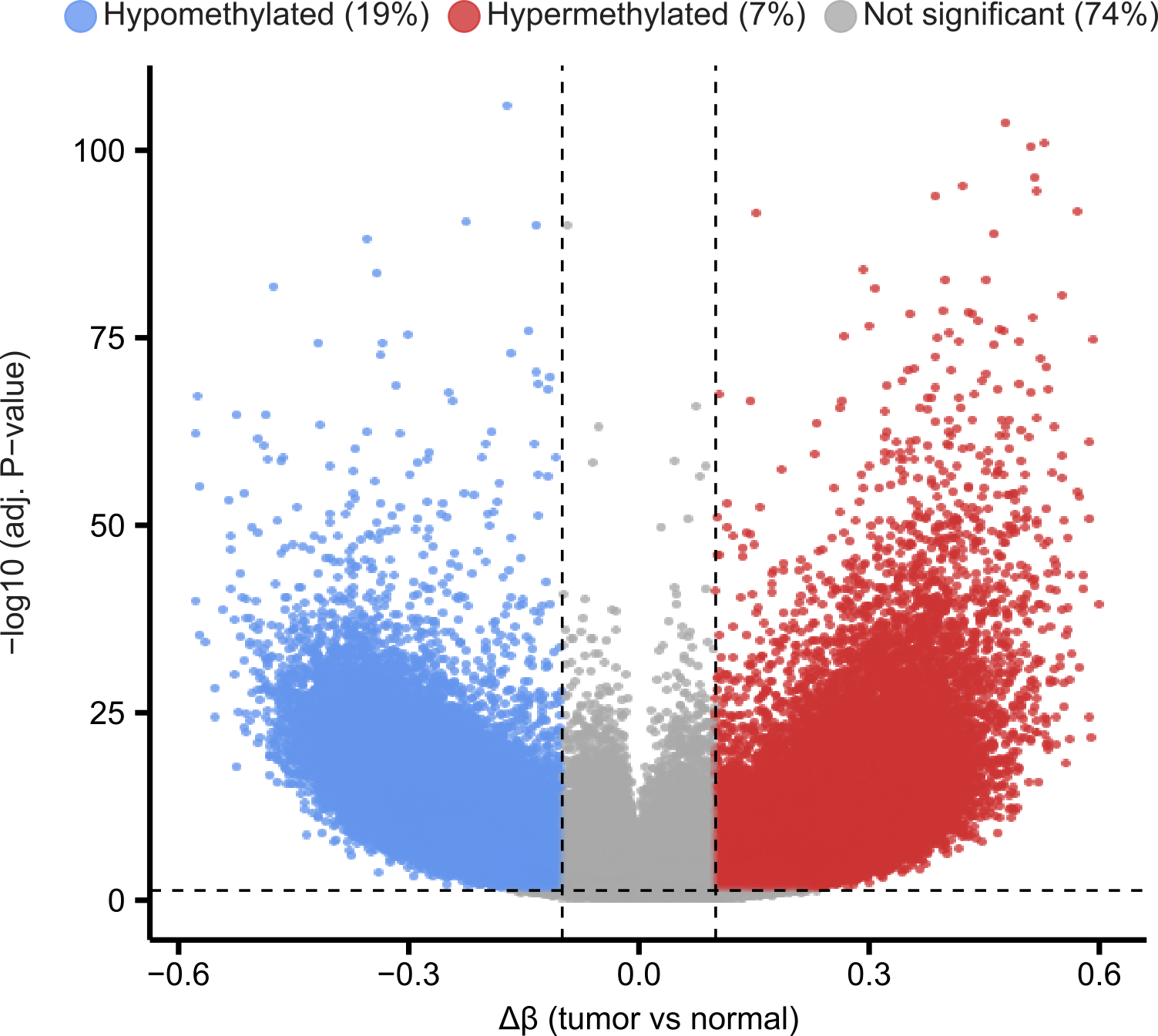
**Figure S4. Differential methylation analysis between tumor and normal samples.** Volcano plot of probe-wise differential methylation analysis comparing tumor (*n* = 274) and normal (*n* = 19) samples. Analysis was performed using linear models implemented in the *limma* package on M-values derived from EPIC array data. The horizontal dashed line indicates the significance threshold corresponding to an adjusted *P* value < 0.05. Vertical dashed lines indicate effect size thresholds for hypermethylation (Δβ > 0.1; red) and hypomethylation (Δβ < -0.1; blue).

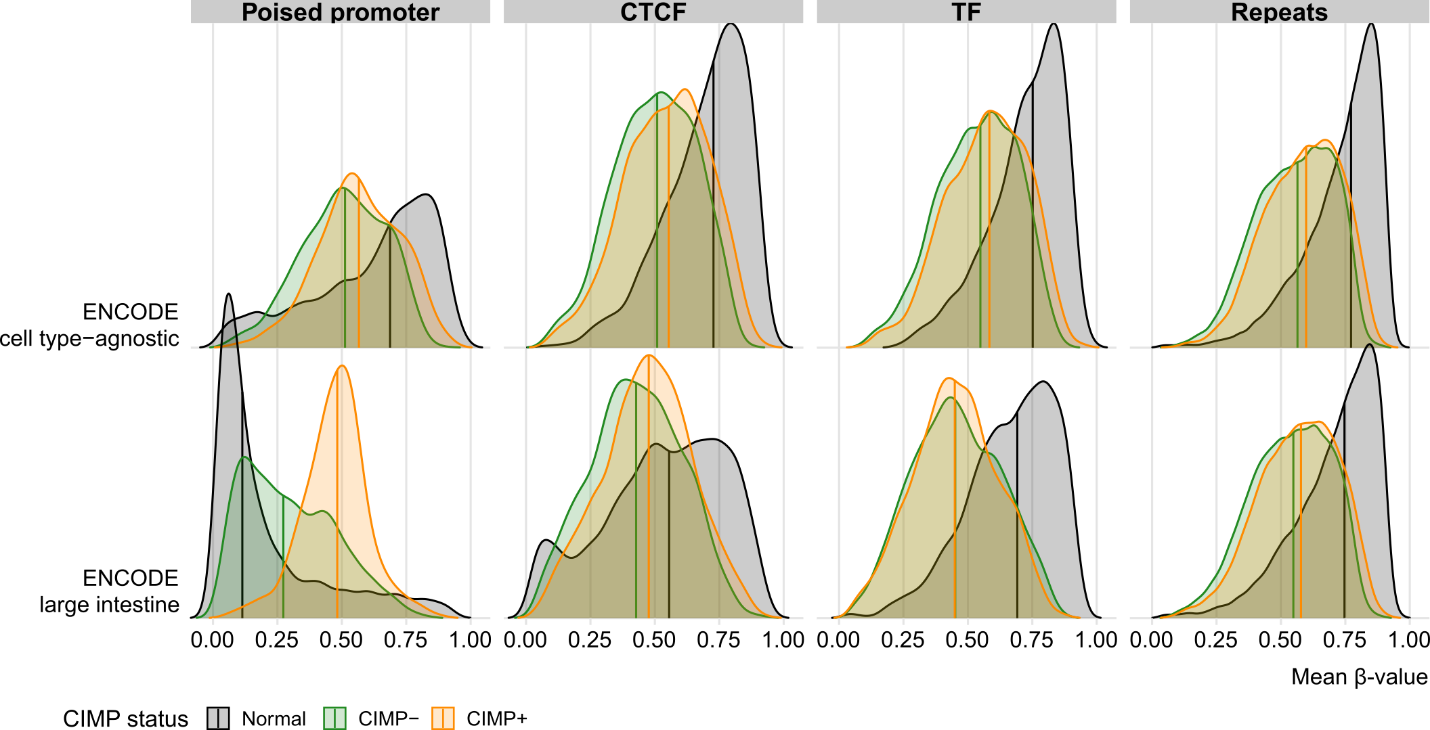
**Figure S5. Distribution of mean β-values across ENCODE-derived probe annotations.** Density ridge plots of average beta values stratified by selected ENCODE-derived probe annotation categories. Tumor samples are colored by CIMP status (green, CIMP-; orange, CIMP+), with normal samples shown in black for reference. Vertical lines indicate median values.

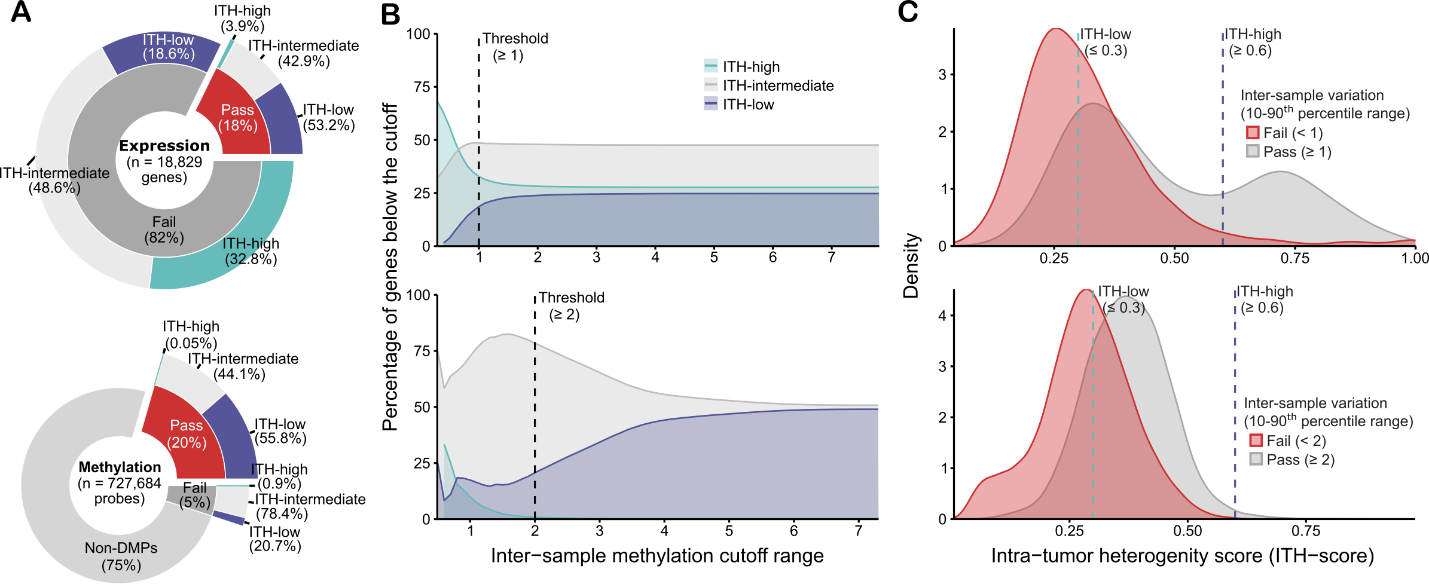
**Figure S6. Distribution of genes and probes by intra-tumor heterogeneity (ITH). (A)** Proportion of genes and probes categorized as ITH-low (≤ 0.3), ITH-intermediate (0.3-0.6), or ITH-high (≥ 0.6) in the multiregional primary tumor cohort (*n* = 274 samples). Non-differential methylated probes (non-DMPs) were not assigned to ITH categories. **(B)** Distribution inter-sample variability, defined using the 10^th^-90^th^ percentile range, stratified by ITH scores. Dashed vertical lines indicate threshold criteria (pass: ≥ threshold; fail: < threshold). **(C)** Density plots of ITH-scores are grouped according to the inter-sample variability filter. Dashed vertical lines indicate the threshold used to define ITH categories.

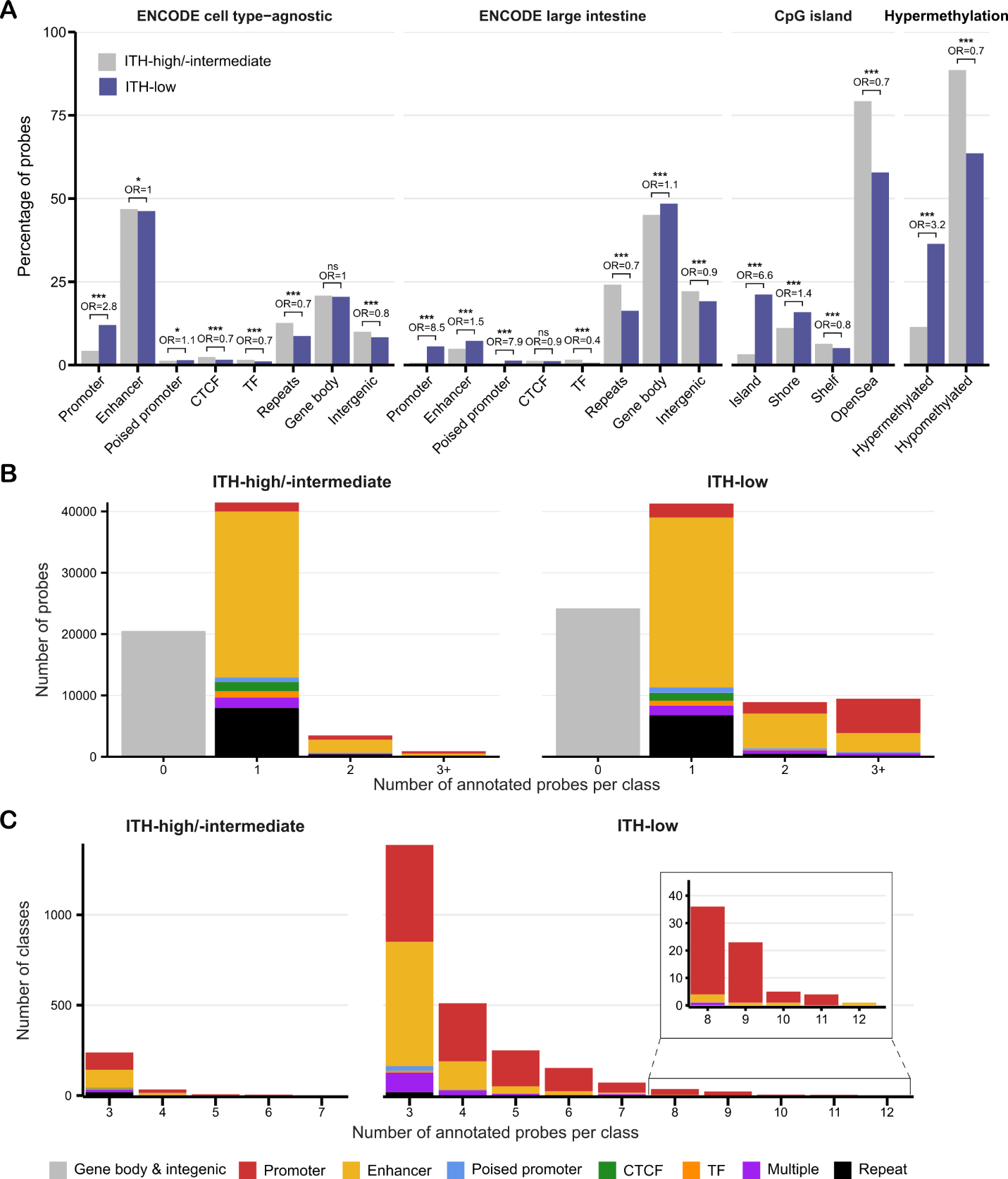
**Figure S7. Distribution of probes categories by intra-tumor heterogeneity (ITH). (A)** Bar plots showing the percentage of probes stratified by ENCODE-derived annotations (cell type-agnostic and large intestine-specific), CpG island context and hypermethylation status. Probes are grouped and colored by ITH category (ITH-high/-intermediate, grey; ITH-high; purple). Stars indicate FDR-adjusted chi-square significance (ns: not significant, *FDR < 0.05, **FDR < 0.01, ***FDR < 0.001) and OR represents the odd ratio between ITH-low and ITH-high/-intermediate. **(B)** Stacked bar plots showing the number of probes mapped to candidate cis-regulatory elements (cCREs), stratified by the cCRE annotation category. The bar with x-axis named “0” shows the number of probes not mapped to any cCRE. Category named “multiple” denotes cCREs assigned to two distinct annotation categories between ENCODE cell type-agnostic and large intestine-specific datasets. **(C)** Stacked bar plots showing the number of cCREs by the number of probes mapped per cCRE (3-12 probes) stratified and colored by cCRE category for each ITH group.

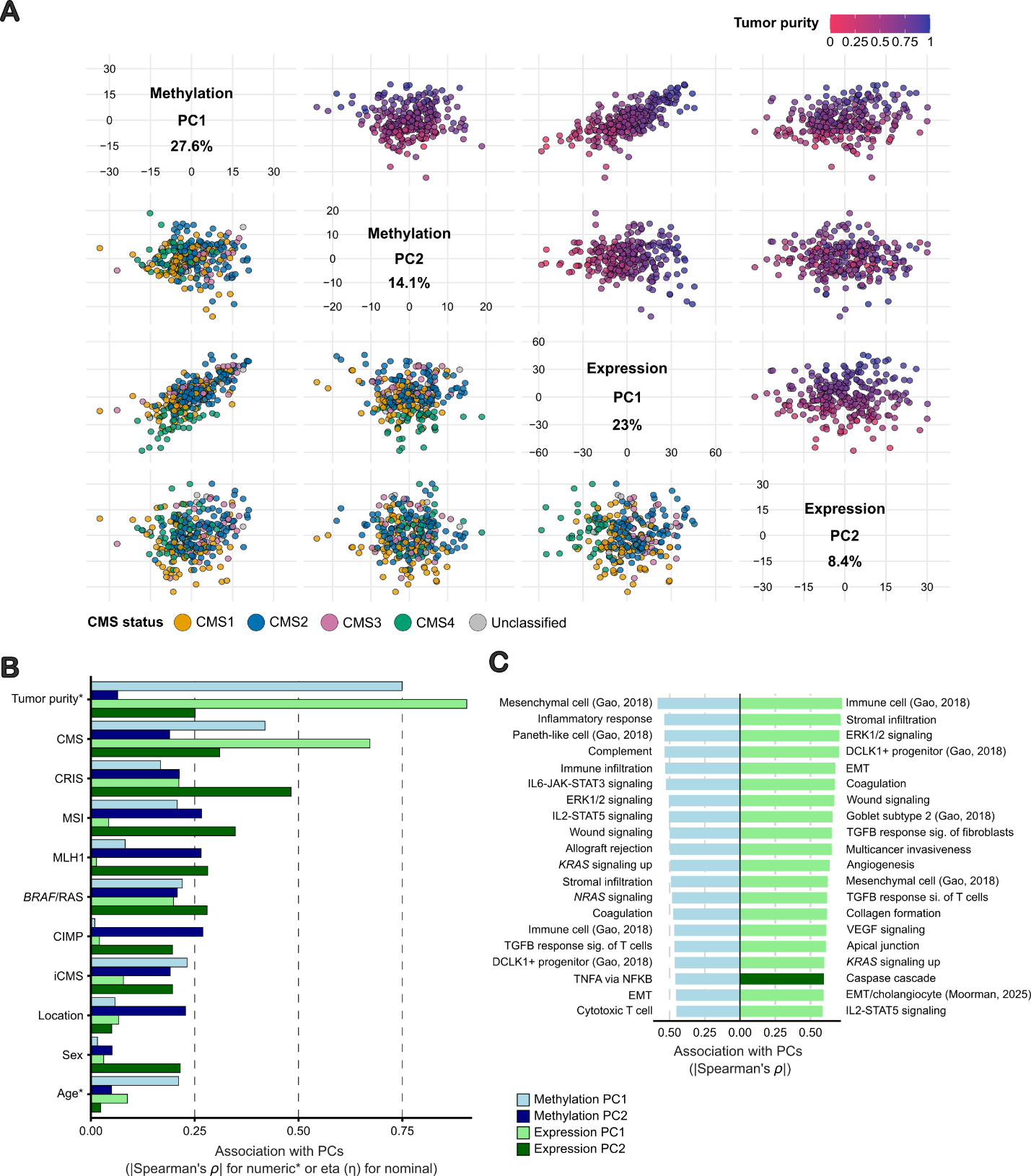
**Figure S8. Comparison of ITH-high/intermediate expression and methylation data. (A)** Scatter plot matrix (SPLOM) of principal component analysis (PCA) results (PC1 and PC2) for each dataset. For methylation, PCA was performed on M-values restricted to ENCODE cCREs with at least three ITH-high/intermediate probes annotated, using the median β-value per cCRE. Percentages indicate the variance explained ratio by each PC in the respective dataset. **(B)** Associations between PCs with molecular and clinicopathological characteristics, and **(C)** the top 20 gene set variation analysis scores. Associations with continuous variables were assessed using Spearman correlation, and associations with categorical variables using correlation ratio eta (η). Bars are colored by PC.

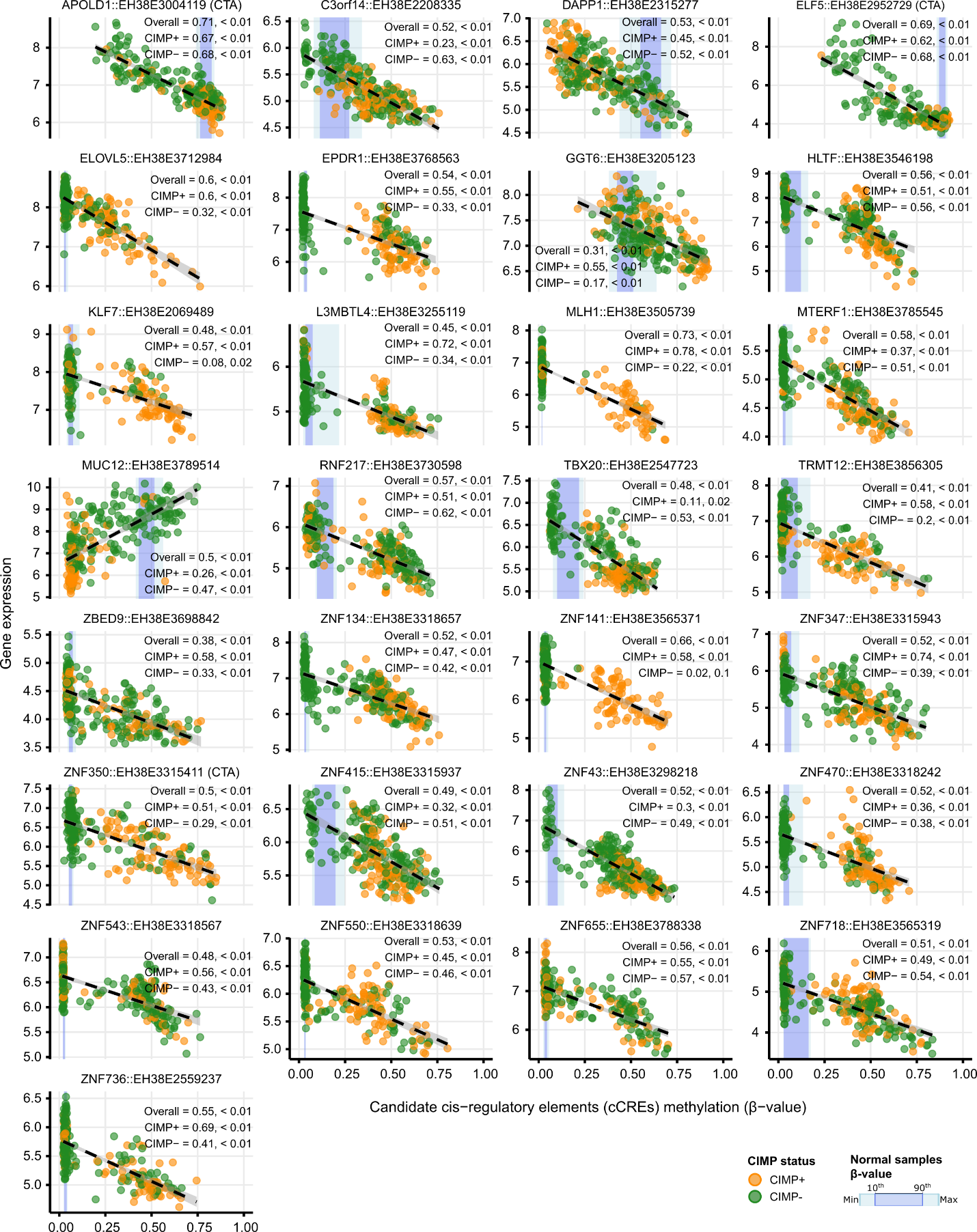
**Figure S9. Main cCRE-gene methylation-expression pairs among ITH-low features.** Scatter plots of the main significantly correlated candidate cis-regulatory elements (cCREs) methylation-expression associations. Models were fitted using linear mixed-effects regression, and marginal R^2^ and *P* values are reported for all samples and stratified by CIMP status. Points are colored by CIMP status. Blue shaded regions indicate the methylation range in normal samples (minimum-maximum and 10^th^-90^th^ percentiles). Black dashed lines represent linear model fits with 95% confidence intervals.

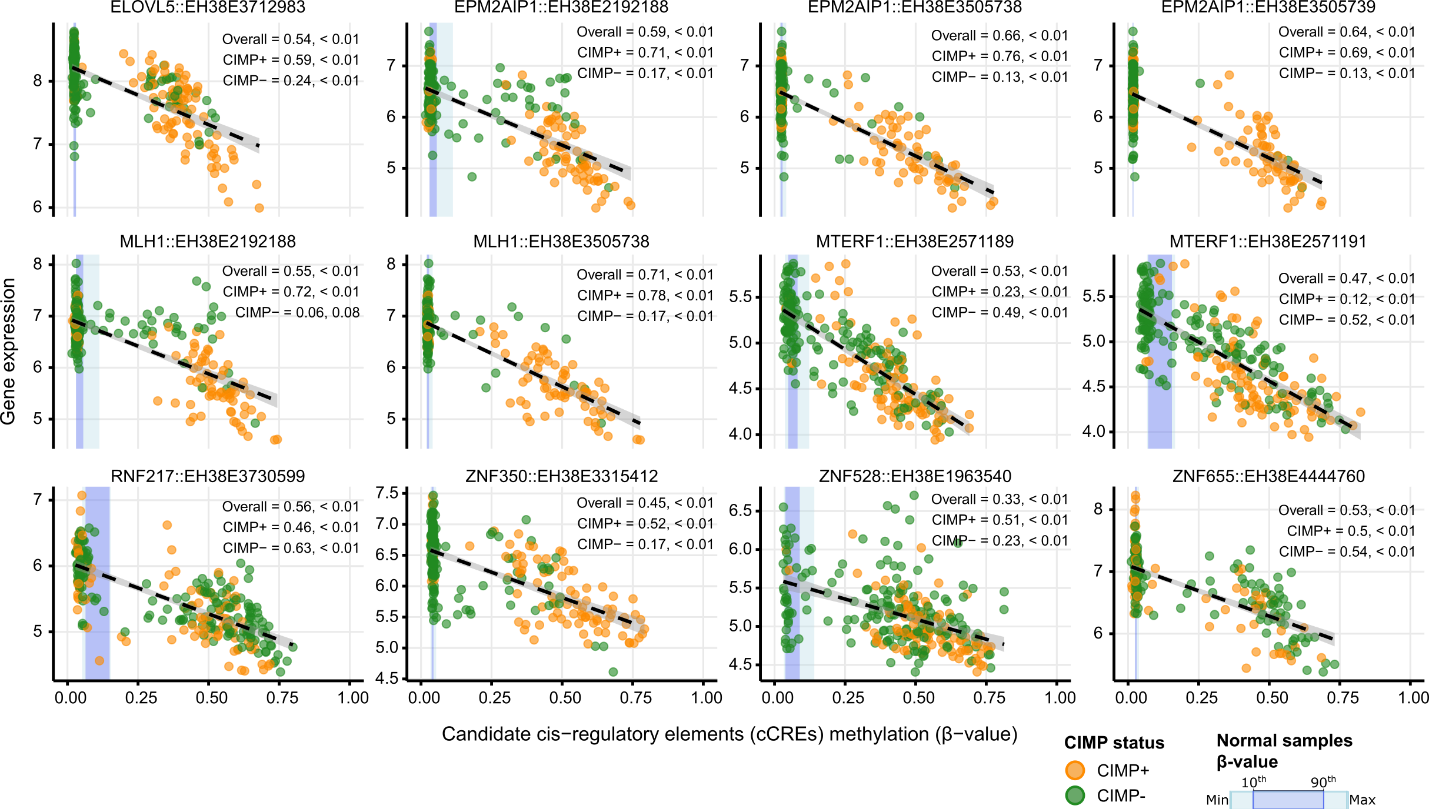
**Figure S10. Secondary cCRE-gene methylation-expression pairs among ITH-low features.** Scatter plots of additional cCRE-gene methylation-expression associations not classified as main. Models were fitted using linear mixed-effects regression, and marginal R^2^ and *P* values are reported for all samples and stratified by CIMP status. Points are colored by CIMP status. Blue shaded regions indicate the methylation range in normal samples (minimum-maximum and 10^th^-90^th^ percentiles). Black dashed lines represent linear model fits with 95% confidence intervals.

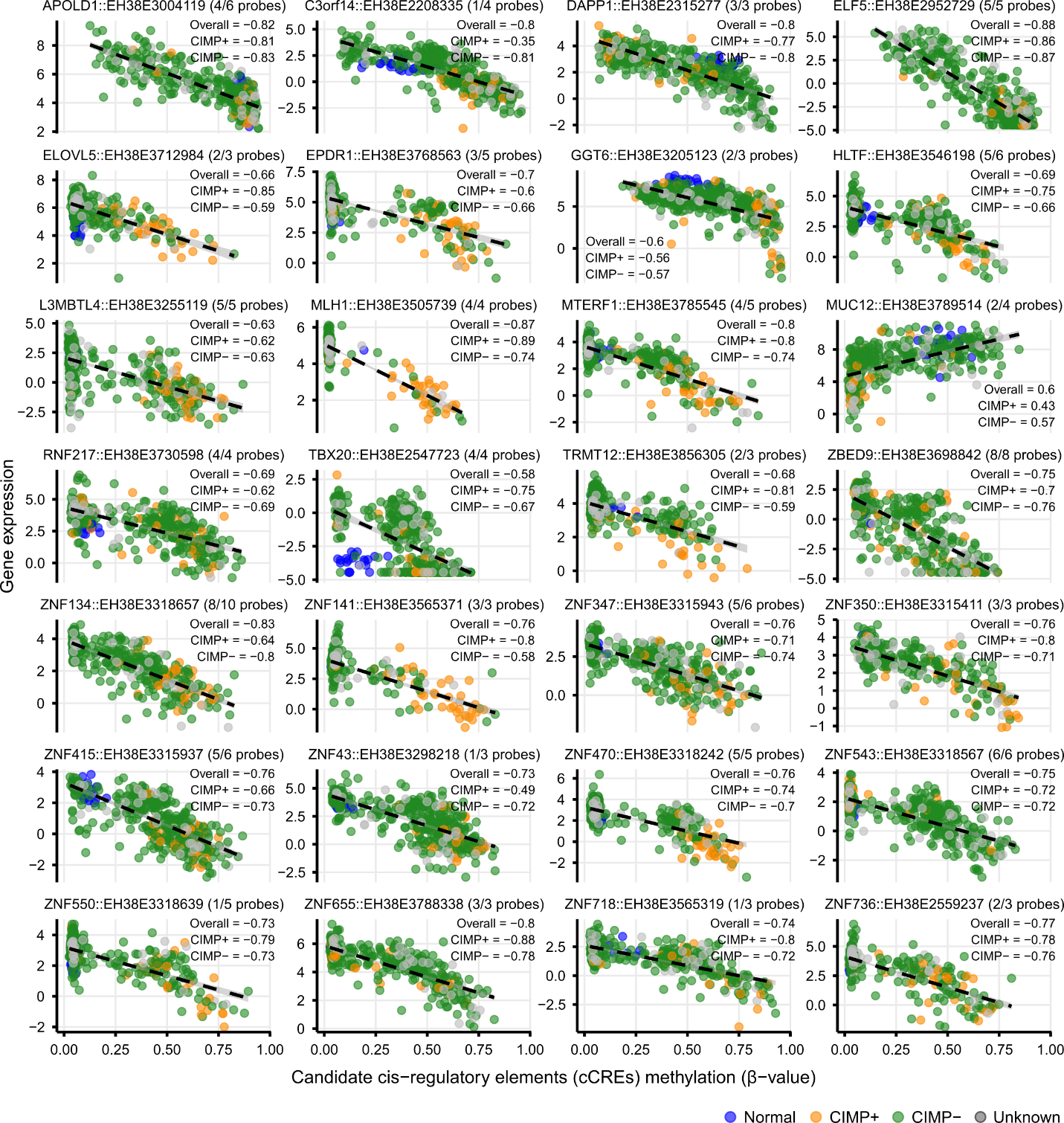
**Figure S11. Validation of main cCRE-gene methylation-expression pairs in the TCGA cohort.** Scatter plots of main cCRE-gene methylation-expression associations evaluated in the TCGA COAD and READ samples with matched methylation and expression data. *KLF7* was excluded due to the absence of corresponding probes in the TCGA dataset. Pearson correlation coefficients are reported for all samples and stratified by CIMP status. Points are colored by sample type (normal) and tumor CIMP status (CIMP+, CIMP-, or unknown). Black dashed lines represent linear model fits with 95% confidence intervals.

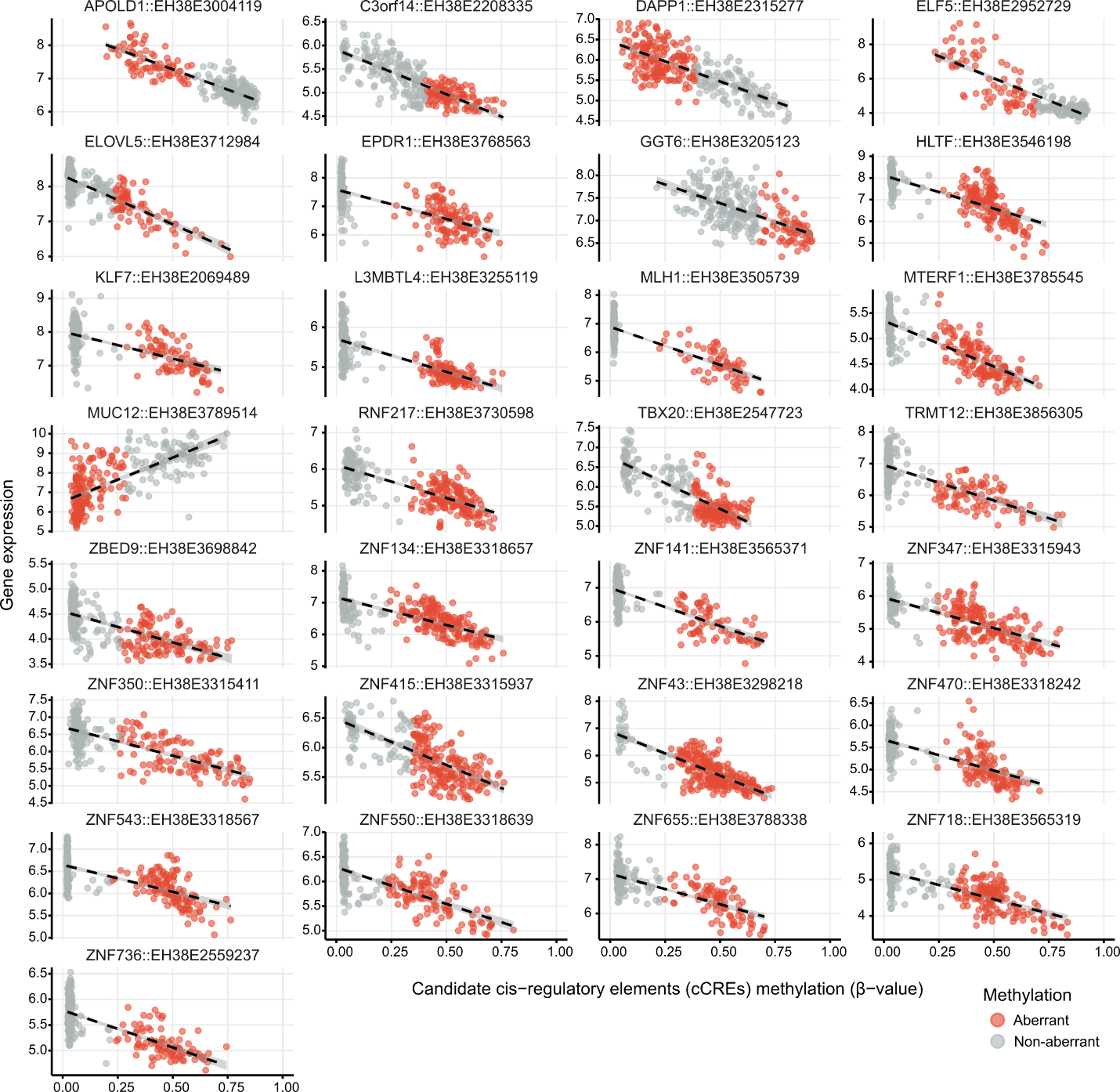
**Figure S12. Definition of aberrantly methylated samples from significant cCRE-gene pairs.** For significant main cCRE-gene correlations, tumor samples were classified as aberrant methylated (red points) if they showed both an absolute Z score > 3 relative to normal samples and an absolute Δβ value > 0.15 compared to the mean methylation of normal samples. Z scores were calculated using the mean and standard deviation of methylation levels in normal samples for each regulatory element.

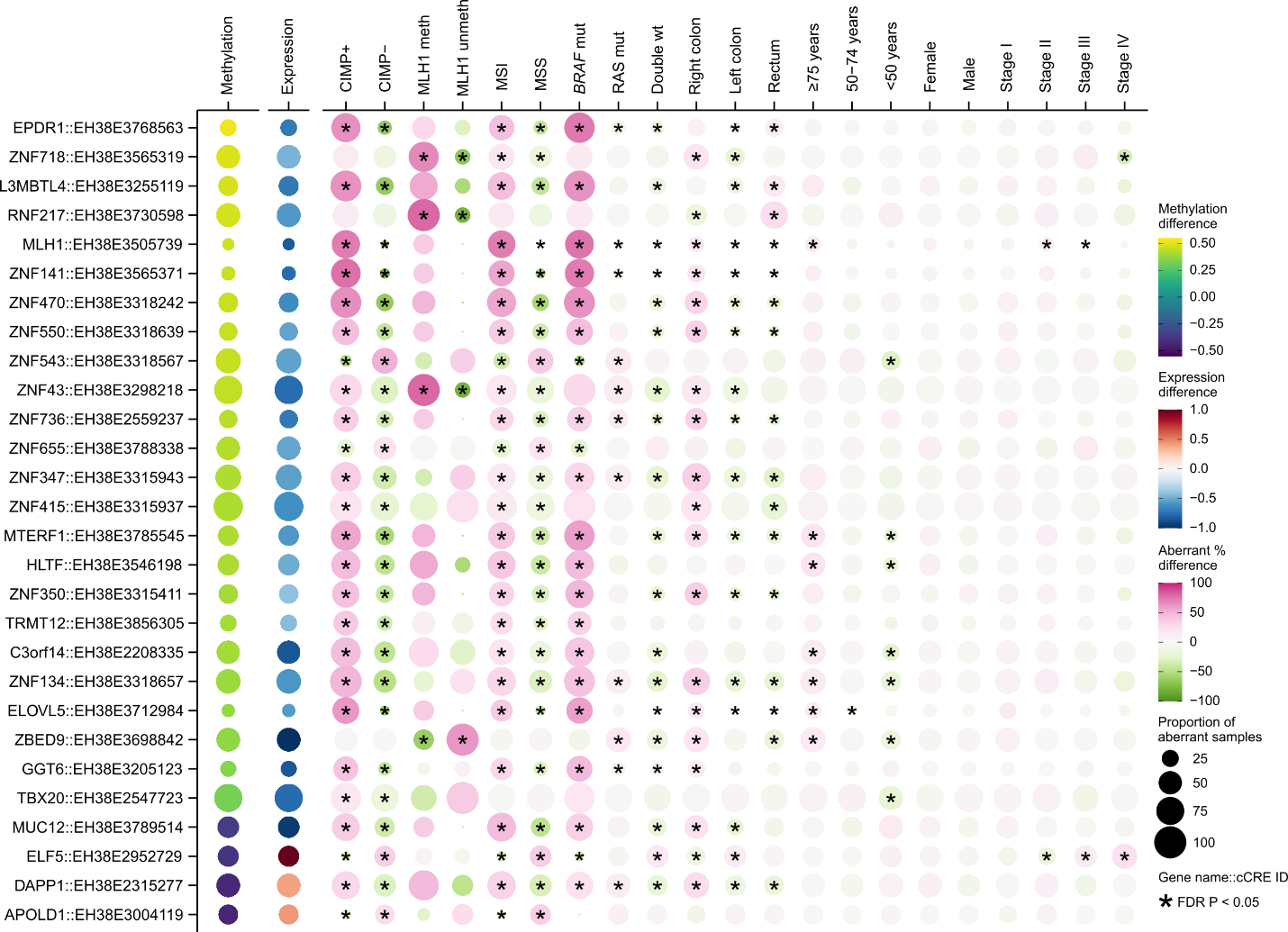
**Figure S13. Validation of molecular and clinicopathological associations of cCRE-gene methylation-expression pairs in the TCGA cohort.** Bubble plot summarizes significant cCRE-gene associations in TCGA COAD and READ samples, showing differences in methylation and gene expression between aberrant and non-aberrant samples, as well as associations with other molecular and clinicopathological characteristics. For molecular and clinicopathological variables, bubble size represents the proportion of samples classified as aberrant within each category, and color represents the aberrant percentage-point difference between the selected variable and all other variables within each category. Associations were tested using logistic regression with FDR correction. Asterisks indicate FDR < 0.05.

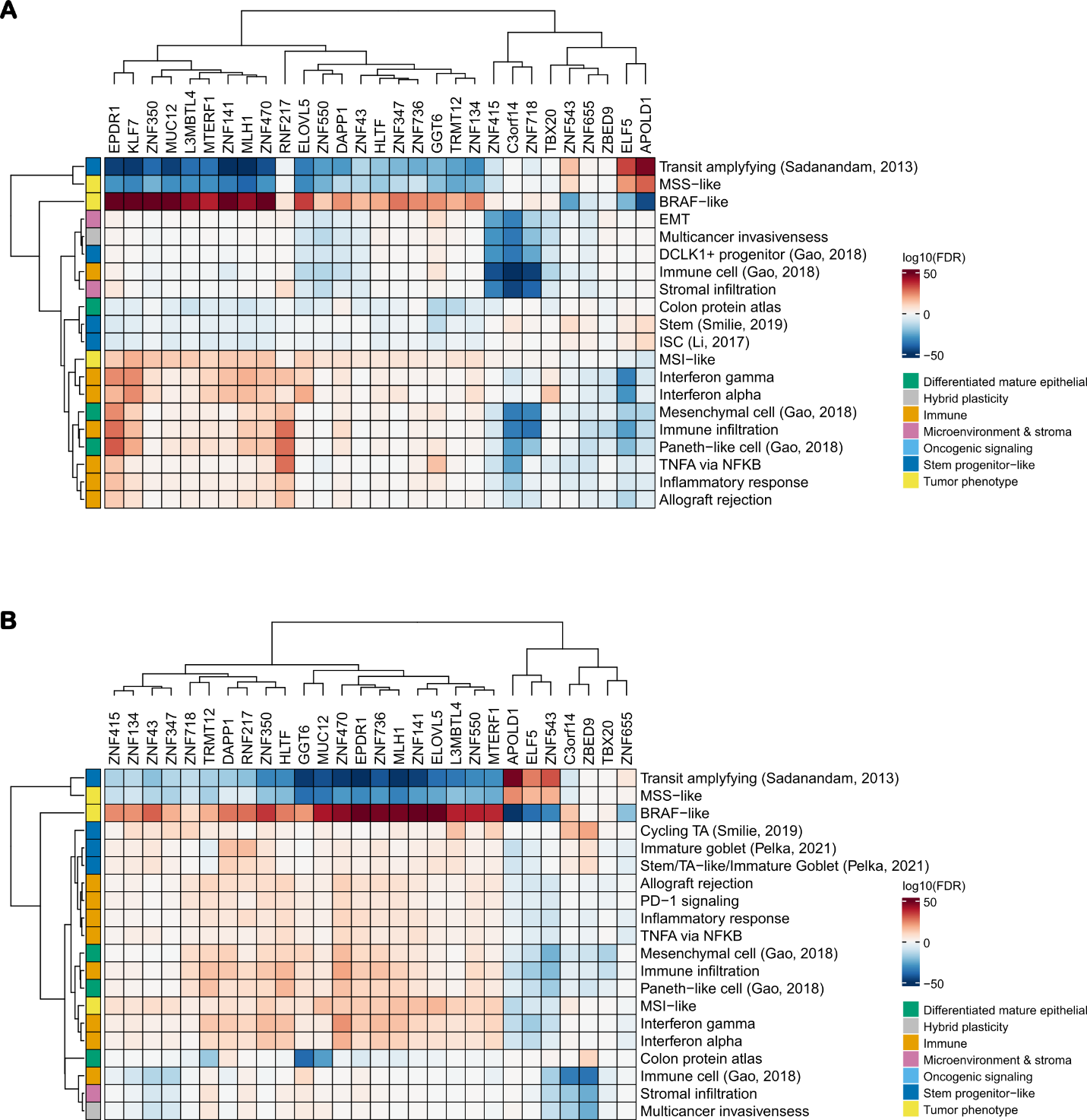
**Figure S14. Validation of gene set enrichments by gene aberrant status across cohorts.** Heatmaps of gene set enrichment analysis results for **(A)** the in-house multiregional cohort and **(B)** the TCGA validation cohort. Color indicates -log10(FDR) for the aberrant versus non-aberrant tumors for each gene. The top 20 pathways were selected based on the mean absolute -log10(FDR) across comparisons. Rows and columns were hierarchy clustered using Euclidean distance and complete linkage.

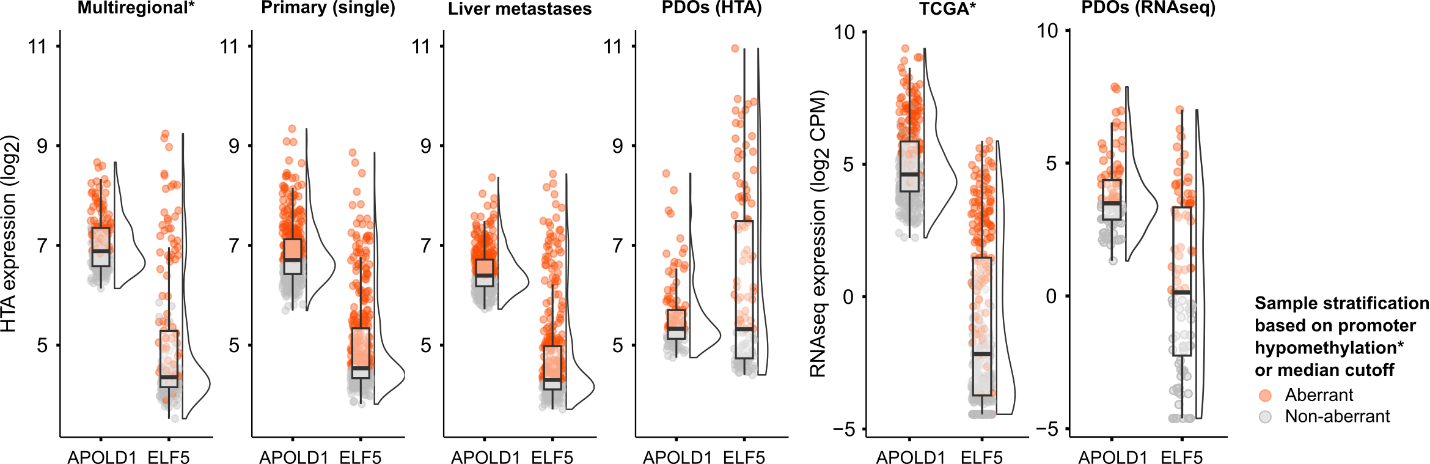
**Figure S15. *APOLD1* and *ELF5* expression across CIMP- and/or MSS samples in the multiple cohorts.** Raincloud plots showing *APOLD1* and *ELF5* expression in CIMP- and/or MSS samples for cohorts profiled with microarrays (in-house multiregional, extended single-primary, liver metastases, and patient-derived organoids cohorts) and RNA sequencing (TCGA and PDOs cohorts) platforms. Aberrant status was defined based on promoter methylation in cohorts with methylation data (in-house multiregional and TCGA series). For cohorts with expression data only, aberrant status was defined using each cohort-specific median expression cutoff for each gene.

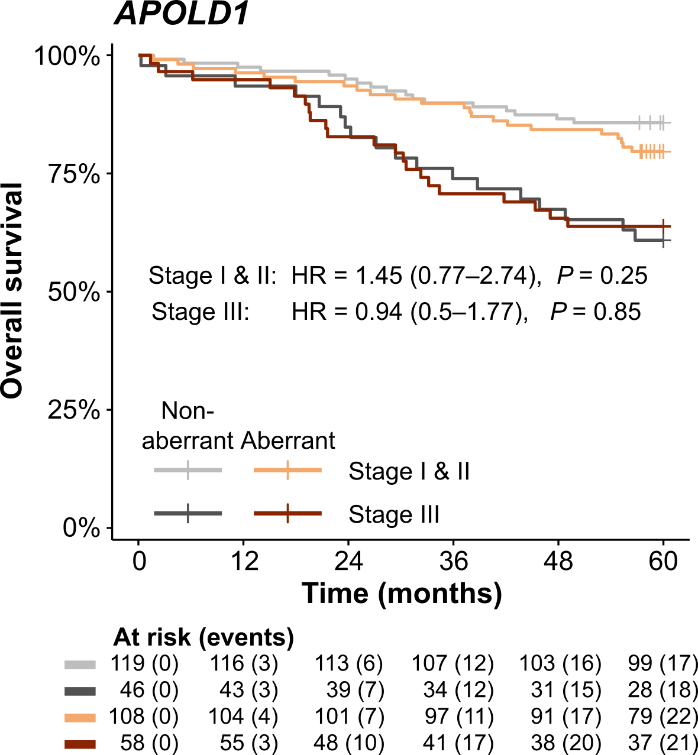
**Figure S16. Survival analysis of *APOLD1* aberrant status in CIMP- and/or MSS CRC.** Kaplan-Meier survival curves of overall survival stratified by *APOLD1* aberrant status and tumor stage (I-II versus III) in CIMP- and/or MSS CRC samples, from the multiregional and extended in-house series (n = 331). Hazard ratios (HRs), 95% confidence intervals, and *P* values are shown for aberrant versus non-aberrant groups.

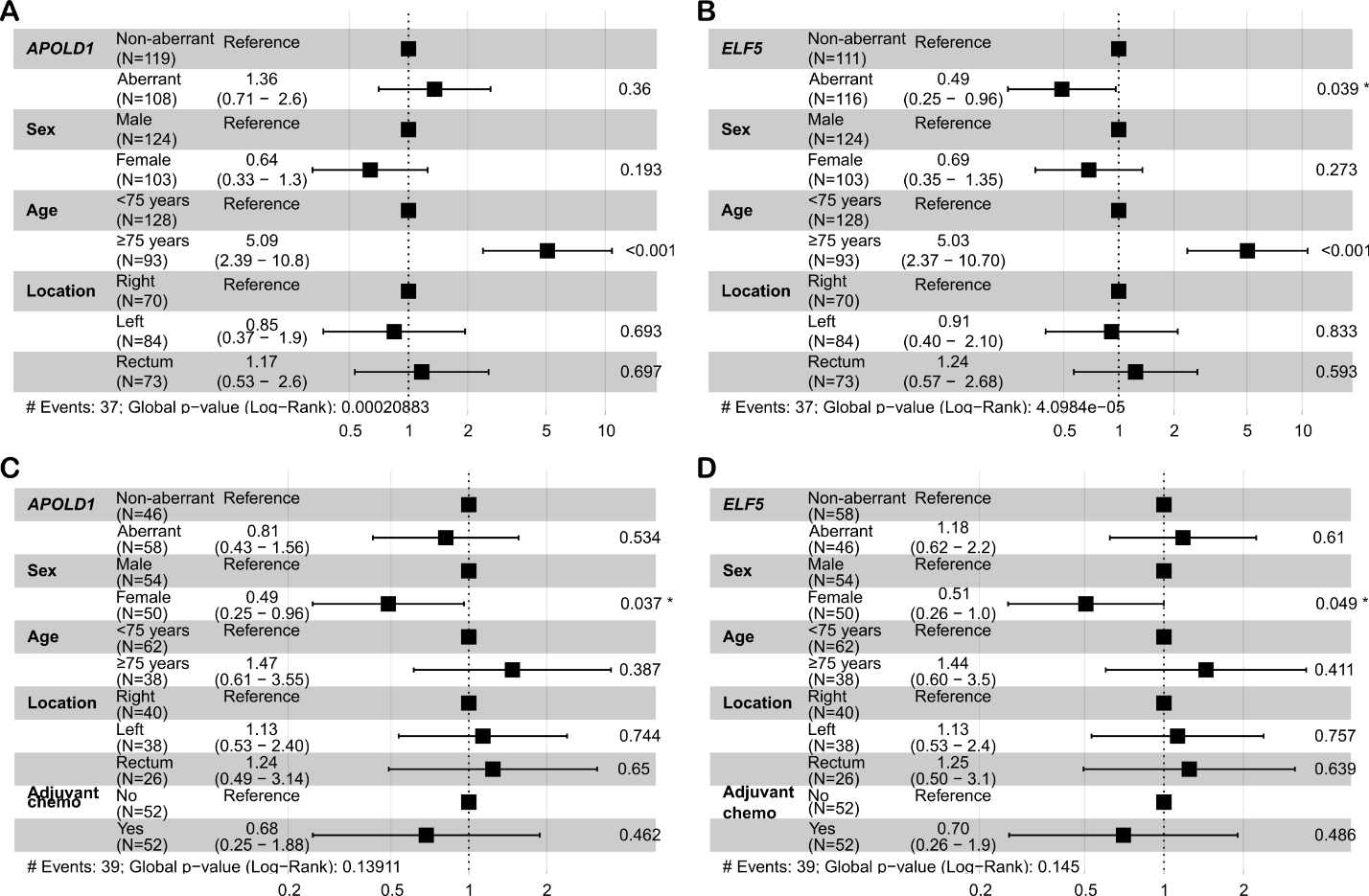
**Figure S17. Clinical characteristics and survival associations of *APOLD1* and *ELF5* aberrant status.** Forest plots for multivariable Cox proportional hazard models with *APOLD1* and *ELF5* aberrant status and clinical characteristics for **(A-B)** stage I-II and **(C-D)** stage III CIMP- and/or MSS tumors.

**Figure S18. *APOLD1* and *ELF5* expression in matched primary colorectal tumors and liver metastases.** Boxplots with paired data points of *APOLD1* and *ELF5* expression in patient-matched primary tumors and liver metastases profiled by HTA arrays (n = 23 patients; 102 samples, all CIMP- and/or MSS). Lines connect paired samples and are colored by the expression difference (metastasis versus primary). *P* values were calculated using paired *t* tests.
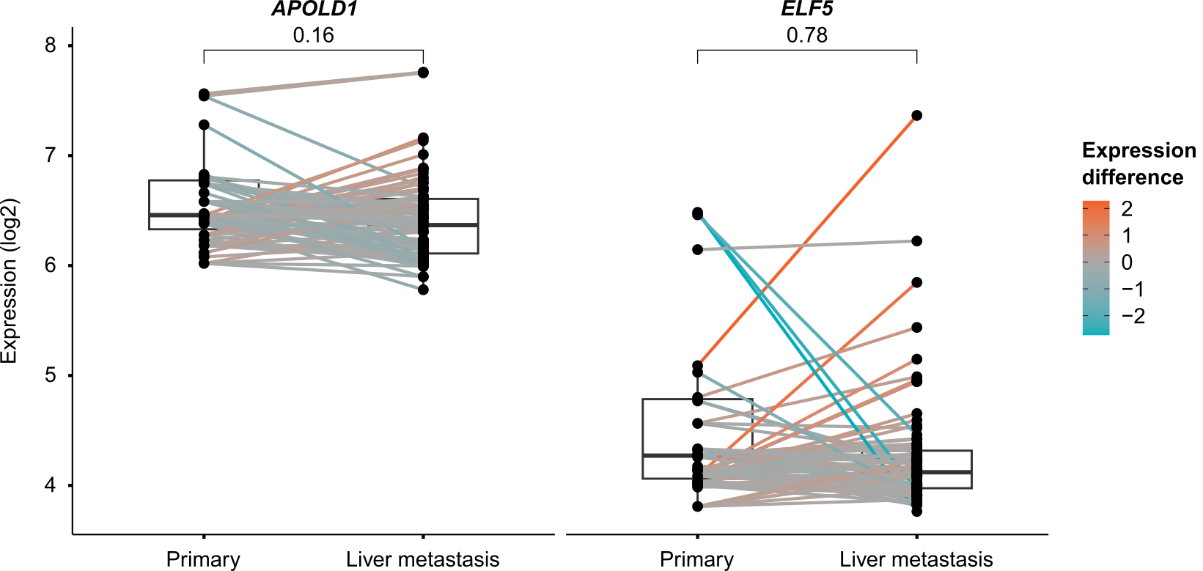
